# Systematic dissection of a complex gut bacterial community

**DOI:** 10.1101/2021.06.15.448618

**Authors:** Alice G. Cheng, Andrés Aranda-Díaz, Sunit Jain, Feiqiao Yu, Mikhail Iakiviak, Xiandong Meng, Allison Weakley, Advait Patil, Anthony L. Shiver, Adam Deutschbauer, Norma Neff, Kerwyn Casey Huang, Michael A. Fischbach

## Abstract

Efforts to model the gut microbiome have yielded important insights into the mechanisms of interspecies interactions, the impact of priority effects on ecosystem dynamics, and the role of diet and nutrient availability in determining community composition. However, the model communities studied to date have been defined or complex but not both, limiting their utility. Here, we construct a defined community of 104 bacterial strains composed of the most common taxa from the human gut microbiota. By propagating this community in growth media missing one amino acid at a time, we show that branched-chain amino acids have an outsize impact on community structure and identify a pathway in *Clostridium sporogenes* for generating ATP from arginine. We constructed and propagated the complete set of single-strain dropout communities, revealing a sparse network of strain-strain interactions including a novel interaction between *C. sporogenes* and *Lactococcus lactis* driven by metabolism. This work forms a foundation for studying strain-strain and strain-nutrient interactions in highly complex defined communities, and it provides a starting point for interrogating the rules of synthetic ecology at the 100+ strain scale.

## INTRODUCTION

Model systems have proven invaluable for the development of mechanistic insight in biology (Müller and Grossniklaus, 2010). Although much has been learned from detailed studies of individual gut commensal species (Cullen et al., 2015; Cuskin et al., 2015; Sonnenburg et al., 2010; Wexler and Goodman, 2017), models of the gut microbiome are less well developed (Blasche et al., 2017; Pacheco and Segrè, 2019; Walter et al., 2018; Widder et al., 2016; Xavier, 2011). Pioneering efforts showed that a synthetic community can model the impact of diet on the microbiome (Faith et al., 2011), identified genes required for *Bacteroides thetaiotaomicron* growth in the mouse intestine in the presence of a 15-member community (Goodman et al., 2009), and demonstrated that complex communities composed of species isolated from a single donor can stably colonize mice (Goodman et al., 2011). More recently, defined communities of up to 20 naturally occurring (Abreu et al., 2019; Friedman et al., 2017; Gutiérrez and Garrido, 2019; Hoek et al., 2016; Medlock et al., 2018; Patnode et al., 2019; Sanchez-Gorostiaga et al., 2019; Yurtsev et al., 2016) or genetically engineered (Hart et al., 2019; Hsu et al., 2019; Kong et al., 2018; Mee et al., 2014; Ziesack et al., 2019) bacterial strains have been studied *in vitro* or in mice, revealing insights into the mechanisms of interspecies interactions. Complex but undefined communities from the gut, soil, and plants have also been studied in detail, illuminating the role of priority effects and the environment—especially nutrient availability—in determining community composition and dynamics (Aranda-Díaz et al., 2020; Goldford et al., 2018; Martínez et al., 2018).

Although these studies have provided foundational insights into the ecology of the gut microbiota, the synthetic communities used have been defined or complex, but not both. An optimal model system would have both features: Near-native complexity would allow a model microbiome to capture properties of an ecosystem that are missing from simpler model systems, including emergent phenomena such as resilience to perturbation (Dethlefsen and Relman, 2011; Ng et al., 2019) and cooperative metabolism (Morris et al., 2013). Moreover, complex consortia are a promising starting point for *in vivo* studies of the gut microbiome, for which they are better suited to model community-level phenomena such as immune modulation and the formation of structured multispecies biofilms.

Complete definition (i.e., communities composed entirely of known organisms) would enable reductionist experiments to probe mechanism. Studies with relatively simple defined communities have demonstrated the power of strain dropout and gene deletion experiments for probing community function. The ability to construct communities with defined composition is especially relevant in the context of experiments testing whether phenotypes can be transferred to germ-free mice via fecal transplant (Gopalakrishnan et al., 2018; Ridaura et al., 2013; Routy et al., 2018). At present, since transplanted communities are typically undefined, it is difficult to uncover the mechanisms underlying these phenomena. A defined model system of sufficient complexity would enable reductionist follow-up experiments, bringing the gut microbiome in line with other model systems in which mechanistic studies are possible.

Here, echoing efforts focused on the plant microbiota (Bai et al., 2015; Carlström et al., 2019; Lebeis et al., 2015), we constructed a complex defined community that contains the most prevalent bacterial species in the human gut microbiome. We demonstrate that the assembly of this 104-member community is reproducible even for very low abundance species. By systematically perturbing this community and its growth medium, we uncover a set of strain-nutrient and strain-strain (e.g. syntrophic) interactions that underlie its composition. This work constitutes a starting point for studying complex synthetic communities at high resolution across orders of magnitude of relative abundance.

## RESULTS

### Designing and building a complex synthetic community

We set out to design a community consisting of the most common bacterial strains in the human gut microbiome. We analyzed metagenomic sequence data from the NIH Human Microbiome Project (HMP) to determine the most prevalent organisms—those that were present in the largest proportion of subjects, regardless of abundance. Although the HMP is not broadly representative of microbiomes from diverse geographies and ethnicities (Deschasaux et al., 2018; He et al., 2018; Sonnenburg and Sonnenburg, 2019), this data set was well suited to our purposes since it was sequenced at very high depth, enabling us to identify low-abundance organisms that are nevertheless highly prevalent (Kraal et al., 2014). After rank-ordering bacterial strains by prevalence, we found that ∼20% (166/844) were present in >45% of the HMP subjects. Of these 166 strains, we were able to obtain 99 from culture collections or individual laboratories (**Figure 1A**). The profiled strains of three additional species were unavailable, so we used alternative strains of the same species (*Lactococcus lactis* subsp. *lactis* Il1403, *Bacteroides xylanisolvens* DSM 18836, and *Megasphaera sp*. DSM 102144). We added two additional strains to enable downstream experiments: *Ruminococcus bromii* ATCC 27255, a keystone species in polysaccharide utilization (Ze et al., 2012); and *Clostridium sporogenes* ATCC 15579, a model gut *Clostridium* species for which genetic tools are available (Dodd et al., 2017; Funabashi et al., 2020; Guo et al., 2019). Together, these 104 strains resemble the phylogenetic distribution of a typical Western human gut community (**Figure S1**). Notably, unlike other defined communities used to model the gut microbiome, our consortium is within ∼2-fold of the estimated number of strains in a typical human gut (Faith et al., 2013; Qin et al., 2010).

**Figure 1:**
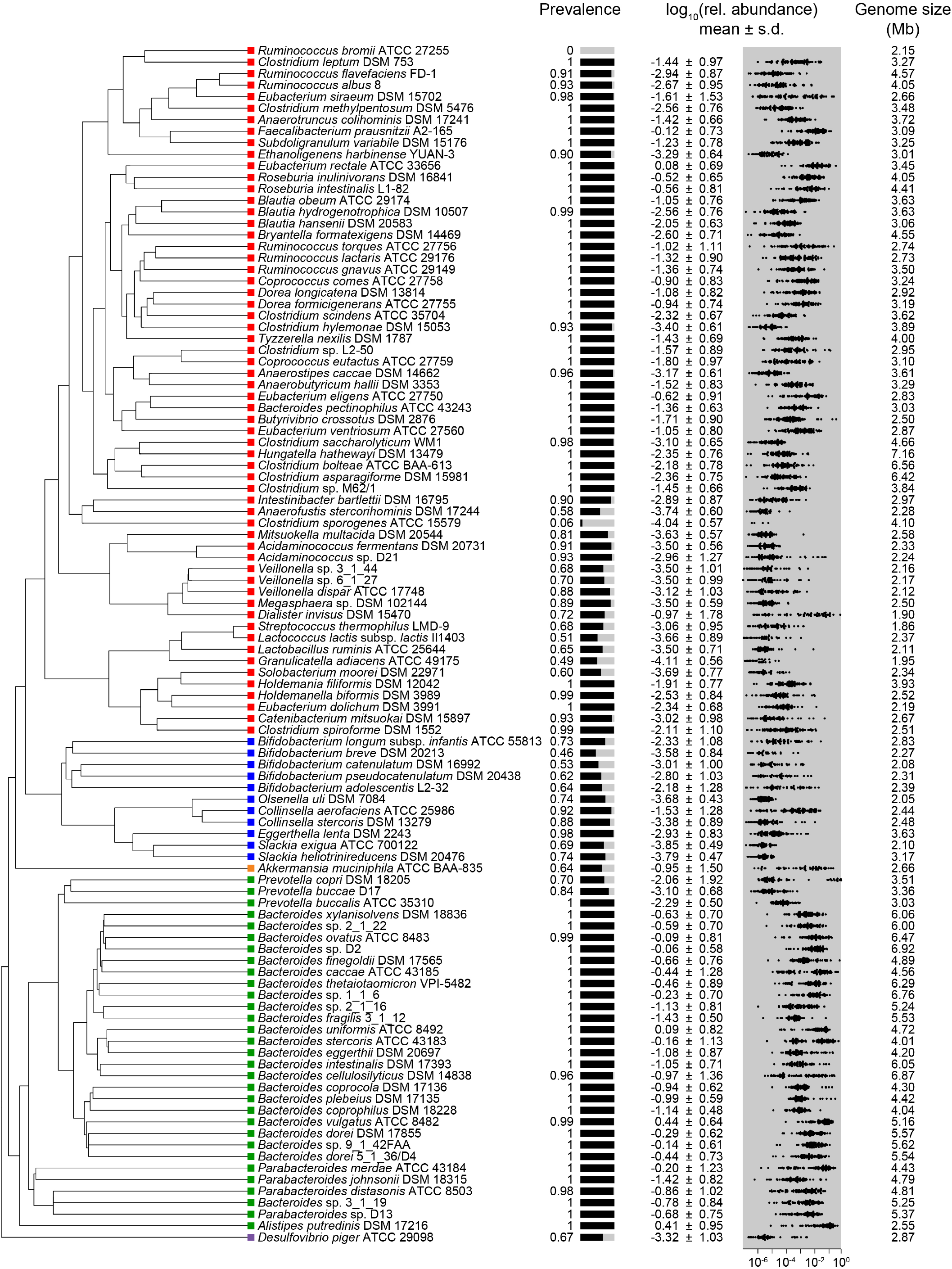
A complex gut bacterial community. A phylogenetic tree of the 104 strains in the community based on a multiple sequence alignment of conserved single-copy genes. The community was designed by identifying the most prevalent strains in sequencing data from the NIH Human Microbiome Project (HMP). Colors indicate the phylum of each strain: Firmicutes = red, Actinobacteria = blue, Verrucomicrobia = orange, Bacteroidetes = green, and Proteobacteria = purple. Also shown are the prevalence and relative abundance of each strain in the data set from the NIH HMP (*n=*81 subjects), and the size of each strain’s genome.

A streamlined strain growth protocol simplified the assembly of the complete community and single-strain dropouts. By testing the growth of each strain in a panel of candidate growth media, we identified two media, at least one of which supports the growth of all 104 strains (**Table S1**). Each culture was passaged daily 2-3 times with dilution into fresh medium. Growth rates, carrying capacities, and time of entry into stationary phase varied widely across strains and media; by passaging fast-growing strains more frequently than slow-growing organisms, we synchronized culture saturation to the extent possible. Before mixing individually cultured strains, we adjusted the volumes of each culture to achieve similar optical densities. We confirmed that these cultures were pure using metagenomic sequencing and high accuracy read mapping, as described in the next section.

### Development of a highly accurate metagenomic read-mapping pipeline

Having assembled a community of 104 strains, we next addressed how to quantify the abundance of each strain accurately, a major challenge in light of our expectation that some strains would be present at low abundance. Various strains in the community have identical 16S hypervariable sequences, ruling out 16S amplicon-based methods. We considered designing a custom amplicon-based pipeline, but such an approach would require the design and validation of new primer sets for future communities, which we wished to avoid. Instead, we sought to use metagenomic sequencing as a means of quantifying community composition.

To test the performance of existing metagenomic analysis tools, we generated three ‘ground truth’ data sets. The first two consisted of simulated reads generated from the assembled genome sequences of each strain: one in which all 104 strains were equally abundant (to test sensitivity and specificity), and another in which strain abundance varied over five orders of magnitude (to test dynamic range). The third set consisted of actual reads derived from sequencing each strain individually using the same protocol on the same sequencing instrument used for subsequent community analyses. This data set allowed us to account for biases introduced by library construction and sequencing.

We found that metagenomic read mappers based on a combination of Bowtie2 (Langmead and Salzberg, 2012) and SAMtools (Li et al., 2009) were sensitive but inaccurate: there was substantial mis-mapping of reads from one strain to others, such that whole-genome sequencing data from an individual strain was often interpreted as having arisen from multiple strains. Read mis-mapping from any abundant strain would therefore create noise that exceeds signal from low-abundance strains, degrading accuracy. In contrast, algorithms that focus on a few universal genes or unique k-mers such as MetaPhlan2 (Truong et al., 2015), MIDAS (Nayfach et al., 2016), Kraken2/Bracken (Lu et al., 2017; Wood et al., 2019), IGGsearch (Nayfach et al., 2019), or Sourmash (Titus Brown and Irber, 2016) were generally accurate to the species level, but since they only use a small fraction of the reads (<1%), their ability to detect low-abundance or closely related strains is limited.

To address these challenges, we developed a new algorithm, NinjaMap (**Figure 2A**). Taking advantage of the fact that every strain in our community has been sequenced, NinjaMap quantifies strain abundances with high accuracy across >6 orders of magnitude. In brief, NinjaMap considers every read from a sample. If a read does not match perfectly to any of the genomes in the community (typically 3-4% of the reads), it is tabulated but not assigned. If a read has a perfect match to only one strain, it is assigned unambiguously to that strain. If a read matches more than one strain perfectly, it is temporarily placed in escrow. After all of the unambiguous assignments are made, an initial estimate of the relative abundance of each strain is computed. Reads in escrow are then fractionally assigned in proportion to the relative abundance of each strain, normalized by the total size of the genomic regions available for unique mapping to avoid bias in favor of strains with large or phylogenetically distinct genome sequences. Finally, relative abundances are computed.

**Figure 2:**
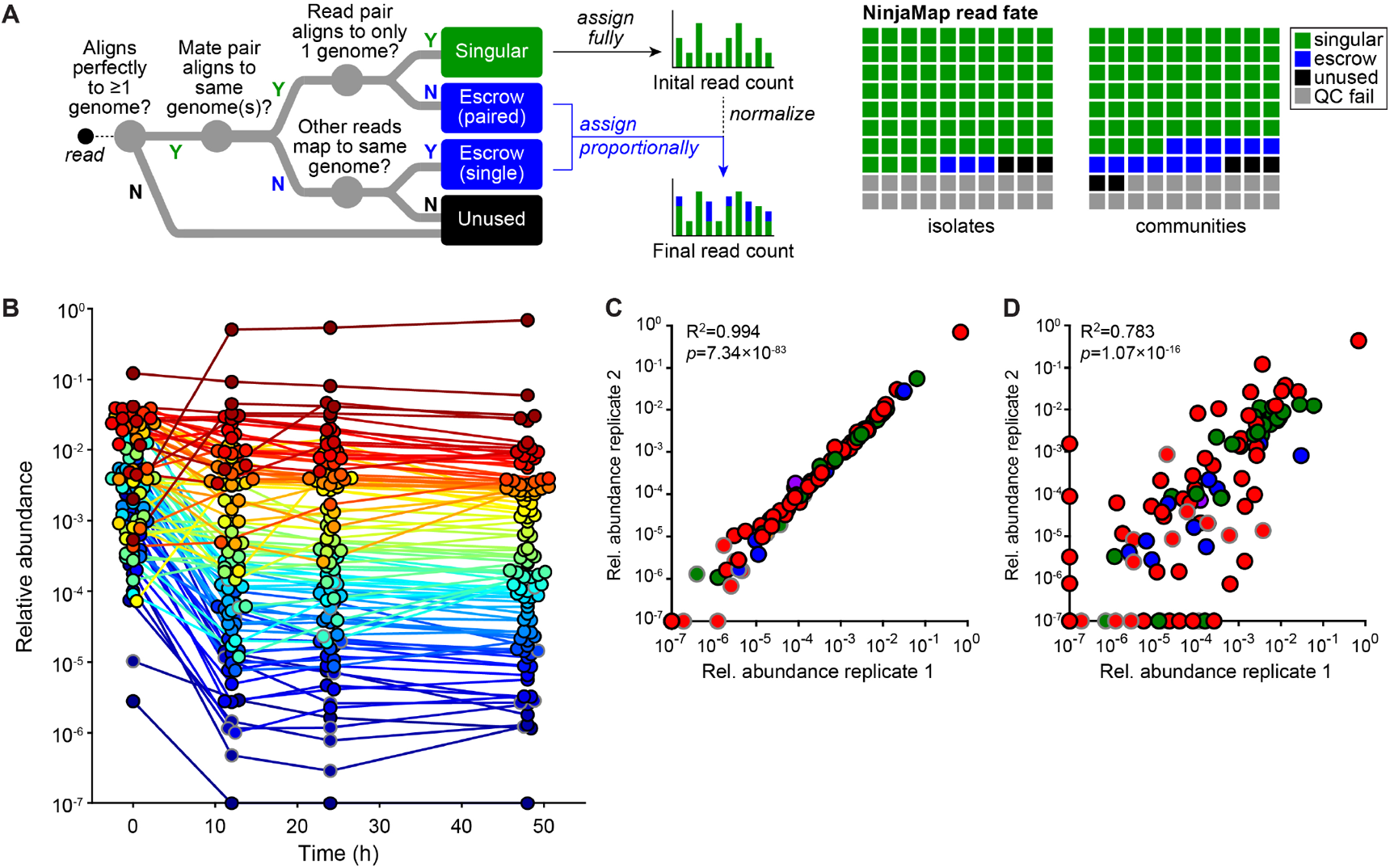
A sensitive and specific read mapping pipeline. (**A**) A schematic of NinjaMap, a new algorithm that quantifies strain abundances in defined communities with high accuracy. Reads that match a single genome unambiguously are assigned to that genome; reads that match multiple genomes are placed in escrow. An initial estimate of the relative abundance of each strain is computed from the unambiguous alignments and used to assign escrow reads proportionally. The final read counts are then normalized to obtain relative abundances. (**B**) The community reaches a stable configuration quickly. Each dot is an individual strain; the collection of dots in a column represents the community at a single timepoint. Strains are colored according to their rank-order abundance in the community at 48 h. By 12 h, The relative abundances of strains in the community span six orders of magnitude and remain largely stable through 48 h. (**C**) Communities generated from the same inoculum (i.e., technical replicates) have a nearly identical composition at 48 h. (**D**) Communities generated from two inocula prepared on different days (i.e., biological replicates) have a similar architecture at 48 h. In (C) and (D), the color of each circle represents the phylum of the corresponding species, and circles with gray outlines represent strains whose presence could be explained by read mis-mapping.

To assess the performance of NinjaMap, we performed two tests. First, we assessed the degree of read mis-mapping from and into each strain’s ledger—e.g., we quantified how many reads from *Bacteroides ovatus* ATCC 8483 were mis-assigned to other strains (which would underestimate its abundance in a community), and how many reads from other strains were mis-assigned to *B. ovatus* (which would overestimate its abundance). For simulated reads, most instances of read mis-mapping resulted in relative abundance errors <10^−4^ (**Figure S2A**). For actual reads, mis-mapping was more frequent but still typically below a threshold of 10^−3^; most mis-mapping arose from deviations between the database genome sequence and the actual sequence of the strain in our collection (**Figure S2B**). In general, if the abundance of a strain in a community was within 10-fold of what would be expected from mis-mapping, we excluded the strain from analyses (**Methods**).

Second, we used Ninjamap to analyze simulated reads from a 104-strain community. We found that this tool can accurately quantify strains with abundances as low as 10^−5^ in the context of a mixed community of known composition (**Figure 2B**). Thus, NinjaMap is capable of quantifying strains accurately over a wide dynamic range of relative abundances.

### Community construction is highly reproducible

Our protocol for assembling communities with >100 members involves several growth passages and liquid transfer steps, and we were concerned that variability in any step of our protocol could make it difficult to interpret results. To address this concern, we measured the degree of reproducibility in community composition data by constructing and propagating the 104-member community multiple times. We included technical replicates to assess variation in bacterial growth, DNA extraction, and sequencing, and biological replicates to determine the impact of differences in the preparation of the inocula. We propagated the communities for 48 h and extracted DNA for sequencing at 0, 12, 24, and 48 h.

Despite our attempts to inoculate equal densities of each strain, the range of densities at *t*=0 spanned several orders of magnitude (**Figure 2B**), with a mean log_10_(relative abundance) of -2.5±0.8 for all detectable strains. Nonetheless, 95/104 strains were detectable at *t*=0; the remaining strains were below the limit of detection or had abundances that could potentially be explained by read mis-mapping. The communities reached a relatively stable configuration by 12 h (**Figure 2B**), with a remarkable degree of reproducibility among biological replicates (**Figure 2C**). Technical replicates were even more similar (**Figure 2D**), indicating that community growth, DNA extraction, and sequencing contributed only modestly to variability. Notably, very low-abundance strains (<10^−4^) were only slightly more variable than high-abundance strains (**Figure 2C**). Taken together, these results indicate that community composition is robust to experimental variation.

### A nutrient drop-out screen to map strain-nutrient interactions in the community

We next sought to explore the network of strain-nutrient interactions in the community. Although much is known about polysaccharide foraging by gut commensals (Martens et al., 2014), far less is known about amino acid utilization, so we performed the experiment in a defined growth medium (SAAC) from which we could remove one amino acid at a time. Since amino acids are often utilized in pairs (Nisman, 1954; Smith and Macfarlane, 1997), eliminating one at a time from a complete background rather than adding one at a time to a null background has a greater potential to reveal phenotypes relevant to community function. Moreover, performing this screen in the context of a complex community (as opposed to the traditional practice of analyzing the growth of isolated strains) enables us to study community-dependent effects such as nutrient competition or mutualism-dependent nutrient utilization.

To map strain-amino acid interactions, we constructed the 104-member community by mixing cultures of each strain propagated in a rich growth medium. We then sub-cultured this consortium in 20 defined growth media, each deficient in a single amino acid; the complete defined medium was used as a control (**Figure 3A**). Samples were taken at 48 h and metagenomic sequencing data were analyzed to determine the impact of amino acid deficiency on the relative abundance of each strain.

**Figure 3:**
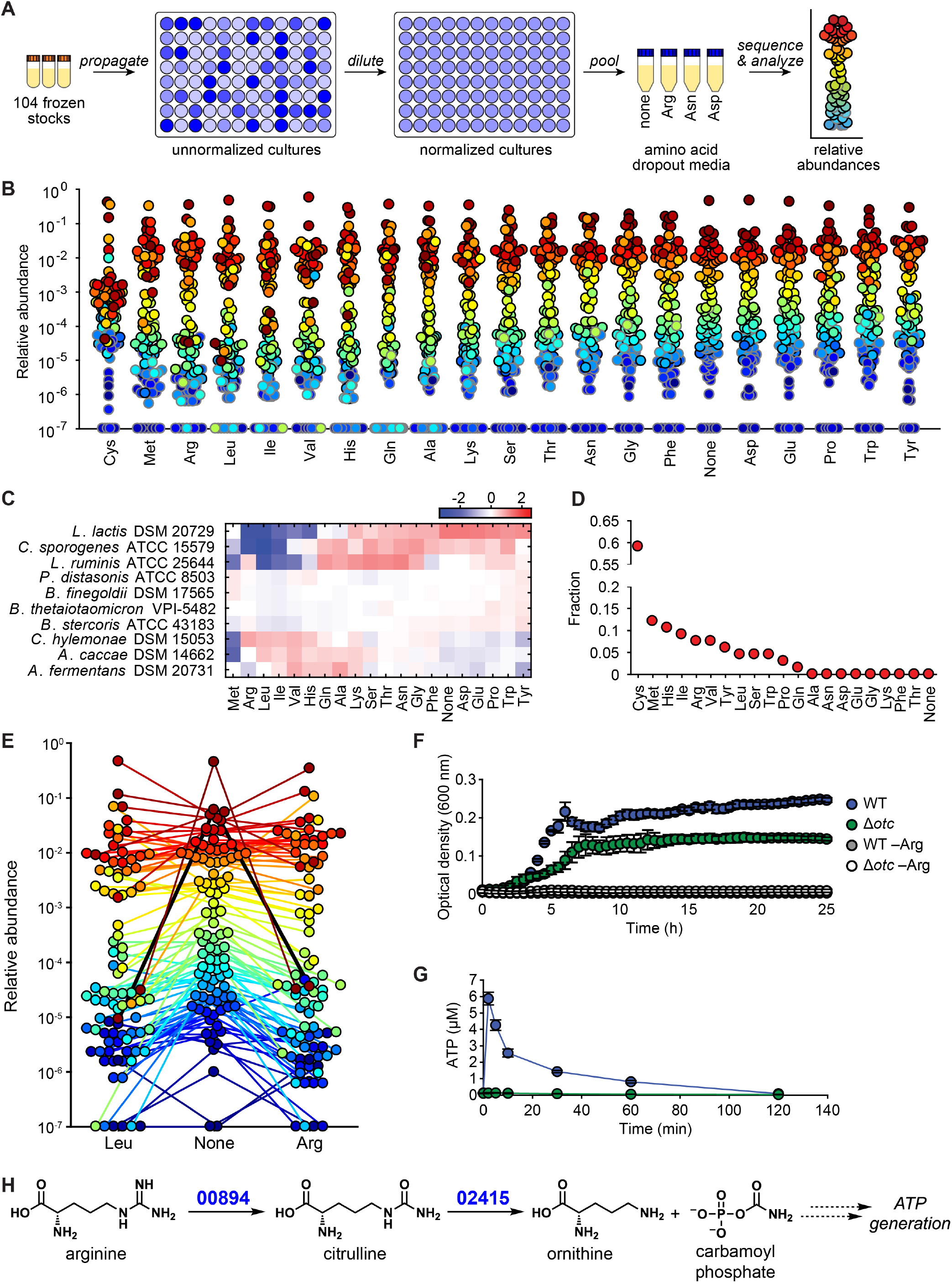
Systematic analysis of strain-amino acid interactions. (**A**) Schematic of the amino acid dropout experiment. Frozen stocks of the 104 strains were used to inoculate cultures that were grown for 24 h, diluted to similar optical densities (to the extent possible), and pooled. The mixed culture was used to inoculate one of twenty defined media lacking one amino acid at a time. After 48 h, communities were sequenced and analyzed by NinjaMap to determine changes relative to growth in the complete defined medium. (**B**) Community composition is impacted by amino acid dropout. Each dot is an individual strain; the collection of dots in a column represents the community at a single timepoint. Strains are colored according to their rank-order abundance in the community grown in complete defined medium. Strains whose relative abundance could be explained by read mis-mapping from a more abundant strain in the same sample are plotted with a gray outline. Undetected strains were set to 10^−7^ for visualization. (**C**) A heat map showing the log_10_(relative abundance) normalized to strain mean for a subset of strains (full set is shown **Figure S3A**). The Firmicutes *L. lactis, C. sporogenes*, and *L. ruminis* grow less robustly in the absence of Leu and Ile. (**D**) The effect of amino acid removal varies widely across amino acids. A *z*-score was calculated based on the standard deviation of strain abundance across all samples except the cysteine dropout. The fraction of strains with |*z*|>2 is shown for each amino acid dropout (*n*=66). (**E**) The absence of leucine or arginine leads to a large decrease in *C. sporogenes* relative abundance. Strains are colored according to their rank-order abundance in the community grown in complete defined medium. Only strains that were detected in at least one of the three samples were included (*n*=92). *C. sporogenes* is highlighted in black. Undetected strains were set to 10^−7^ for visualization. (**F**) *C. sporogenes* growth in complete defined medium is dependent on the presence of arginine, and ornithine transcarbamoylase (*otc*) is partially responsible for Arg metabolism. Wild type *C. sporogenes* and a Δ*otc* mutant were grown in complete defined medium +/- Arg. Growth curves depict the mean of 3 replicates. Error bars represent 1 standard deviation. (**G**) *C. sporogenes* requires *otc* to produce ATP from arginine. Intracellular ATP levels in *C. sporogenes* incubated in PBS containing 2 mM Arg are shown. (**H**) A proposed pathway for Arg metabolism in *C. sporogenes*. Based on these data, we propose that Arg is converted to citrulline by the putative Arg deiminase CLOSPO_00894; citrulline is then hydrolyzed to ornithine and carbamoyl phosphate by the putative ornithine transcarbamoylase CLOSPO_02415, leading to the production of ATP.

### Global analysis of strain-amino acid interactions

To identify strain-amino acid interactions, we tabulated strains whose relative abundance deviated significantly from the mean across conditions, taking advantage of the fact that that most amino acid dropouts had little effect on most strains (**Figure 3B**, Methods). When the community was propagated in the complete defined medium, relative abundances spanned >6 orders of magnitude. 36% of the strains were present at 10^−4^–10^−2^ relative abundance, 8 strains were >10^−2^ and 50 were <10^−4^ (**Figure 3B**). NinjaMap was sensitive to strains with relative abundances as low as 10^−6^, enabling us to quantify the 56% of strains that were below the 10^−3^ limit of detection commonly used for metagenomic analyses (Franzosa et al., 2015). Our system is therefore capable of studying low-abundance microbes, some of which are known to have a large biological impact (Buffie et al., 2015; Funabashi et al., 2020).

To identify significant responses, we calculated the standard deviation of the relative abundance of each strain across experiments and computed *z*-scores (**Figure 3C**). Strain-amino acid interactions that were previously identified in mono-culture studies were also observed in our community format. *Anaerostipes caccae*, whose growth is stimulated by methionine (Soto-Martin et al., 2020), decreased in relative abundance in a community grown in methionine-deficient medium (*z*=-3.48). Likewise, *C*. *sporogenes* was impeded by the absence of leucine (*z*=-2.56), a substrate it oxidatively decarboxylates to isovalerate to generate electrons (Guo et al., 2019). These observations demonstrate that even though >100 strains are competing for the same nutrients, the effects of eliminating one amino acid on the growth of one strain are readily observable in the context of a complex community.

Most strains responded to the removal of ≤4 amino acids. Moreover, relative abundances showed little variability, with a mean standard deviation of log_10_(relative abundance) across strains <0.43. Only three strains, all of which are Firmicutes, were responsive to more amino acids: *Lactococcus lactis* DSM 20729, *Clostridium sporogenes* ATCC 15579, and *Lactobacillus ruminis* ATCC 25644 (**Figure S3, Table S2**). Thus, under these growth conditions, most strains are largely insensitive to amino acid removal while a small minority are highly responsive. We note that the response of a strain to amino acid removal may be direct (e.g. due to utilization for energy) or indirect (e.g. amino acid removal impacts an interacting strain).

In contrast, amino acids varied widely in terms of their impact on community composition (**Figure 3D**). More than half of the strains responded to cysteine removal, likely due to its effect as a reducing agent. More than 5% of the strains responded to methionine, histidine, isoleucine, arginine, valine, and tyrosine, while for eight amino acids there were no significant changes to the community at all (**Figure 3D**). Interestingly, there were large differences among similar amino acids: no strains responded to lysine, while 10.6% and 7.6% of the strains responded to histidine and arginine, respectively. The removal of isoleucine, leucine, and arginine had a particularly large impact on community structure: *C. sporogenes* and *L. lactis*, the two most abundant strains when grown in complete defined medium, each decreased >500-fold in relative abundance when any of these amino acids were removed. Thus, certain amino acids are ‘keystone’ nutrients that play an important role in determining community composition.

### *C. sporogenes* uses arginine to generate ATP

Among the 86 candidate strain-amino acid interactions revealed by our screen, we were particularly intrigued by those involving *C. sporogenes*. Although *C. sporogenes* can oxidize and reduce aromatic amino acids (Dodd et al., 2017), its relative abundance was unaffected by the removal of phenylalanine, tyrosine, or tryptophan (**Figure S4**). In contrast, the removal of leucine, isoleucine, and arginine each had large impact on its fitness in the community. The second strongest phenotype was a decrease in relative abundance in the absence of arginine (**Figure 3E**); although *C. sporogenes* is known to metabolize arginine (Venugopal and Nadkarni, 1977; Wildenauer and Winter, 1986), no impact of arginine on growth or energy metabolism had been observed in prior work. To validate and characterize this interaction, we compared *C. sporogenes* growth in complete defined versus arginine-deficient medium. Although *C. sporogenes* grew well in complete defined medium, it exhibited a large growth defect in the absence of arginine (**Figure 3F**), indicating that this amino acid is an important substrate for growth.

*C. sporogenes* can use other amino acids as substrates to support ATP synthesis (Dodd et al., 2017). Hypothesizing that the same is true for arginine, we incubated wild-type *C. sporogenes* in a culture medium deficient in substrates for ATP synthesis. Upon addition of arginine, intracellular ATP levels rose sharply (**Figure 3G**), indicating that *C. sporogenes* generates ATP (directly or indirectly) from arginine.

To identify the enzymes involved in this process, we parsed the *C. sporogenes* genome for pathways known to capture energy from arginine. This search yielded candidate genes for each of the three steps in the arginine deiminase pathway (**Figure 3H**), which catalyzes the net conversion of arginine to ornithine plus CO_2_ and two equivalents of ammonium, generating one equivalent of ATP (Cunin et al., 1986). Using a method we recently developed to construct scarless deletions in *C. sporogenes* (Guo et al., 2019), we generated strains deficient in the putative arginine deiminase (CLOSPO_00894, Δ*adi*) or ornithine carbamoyltransferase (CLOSPO_02415, Δ*otc*). The Δ*otc* mutant was unable to generate ATP in response to arginine provision, consistent with a role for the arginine deiminase pathway in *C. sporogenes* energy production (**Figure 3G**). In contrast, the Δ*adi* mutant showed no defect in arginine-induced ATP production (**Figure S5A**), suggesting the possibility of an alternative pathway to generate citrulline from arginine. Consistent with these observations, the Δ*otc* mutant (but not the Δ*adi* mutant) was deficient in growth in complete defined medium (**Figures 3F, Figure S5B**). The deficiency was partial, suggesting that an alternative pathway can generate energy from arginine under these conditions. Together, these results show that arginine metabolism by the arginine deiminase pathway contributes directly to the cellular ATP pool, augmenting our understanding of how amino acid metabolic pathways contribute to the fitness of a prominent gut commensal within a complex community.

### A strain drop-out screen to map strain-strain interactions

Next, we sought to map strain-strain interactions within the community. A wide variety of interaction types have been characterized (Little et al., 2008; Shank and Kolter, 2009), including mutualistic or commensal interactions based on nutrient exchange (e.g. syntrophies and secondary fermentation) (Morris et al., 2013) and antagonistic interactions based on antibiotic production or nutrient competition (Zipperer et al., 2016). However, relatively little effort has been applied to characterizing strain-strain interactions systematically; previous efforts have focused on interactions in binary culture (Limoli et al., 2019; Traxler et al., 2013; Vetsigian et al., 2011), where the rules and selective conditions are likely distinct from those in a complex community (Bairey et al., 2016).

To address this gap in knowledge, we constructed all 104 single-strain dropout communities (**Figure 4A**, Methods). These 103-member communities as well as the full community were grown in complete defined medium, and samples were taken at 0 and 48 h to assay growth dynamics over the course of a single passage (**Table S3**). We quantified the relative abundance of each strain by analyzing metagenomic sequencing data using NinjaMap.

**Figure 4:**
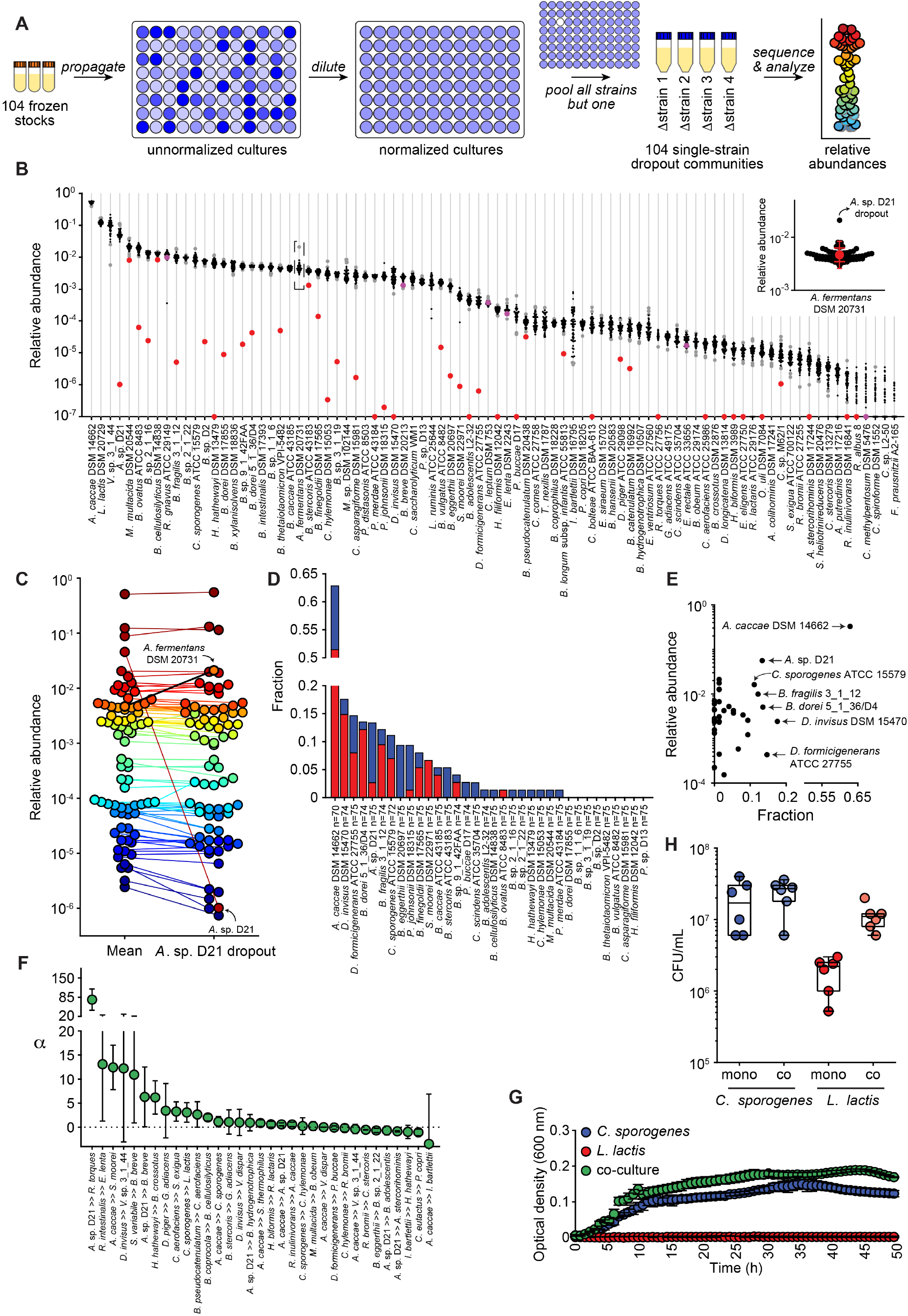
Systematic analysis of strain-strain interactions. (**A**) Schematic of the strain dropout experiment. Frozen stocks of the strains were used to inoculate cultures that were grown for 24 h, diluted to similar optical densities (to the extent possible), and combined into 104 communities, each of which is missing a single strain (i.e., 103-member communities). After 48 h, communities were sequenced and analyzed by NinjaMap to determine changes relative to the growth of the full 104-member community. (**B**) Relative abundances for most strains are narrowly distributed. Each column depicts the relative abundance of an individual strain across a set of 57 strain dropouts (black dots); the relative abundance of each strain in its own dropout is shown as a red dot. Relative abundances in samples with |*z*|>2 are shown as gray dots. Undetected strains were set to 10^−7^ for visualization. Inset: an enlarged view of the relative abundance of *Acidaminococcus fermentans* DSM 20731 across 57 strain dropouts, showing that the elimination of *Acidaminococcus* sp. D21 led to the expansion of *A. fermentans* DSM 20731. The red circle is the mean log_10_(relative abundance); error bars show one (wide bar) or two (narrow bar) standard deviations from the mean. (**C**) Response of the community to the removal of a single strain. Each dot is an individual strain; the collection of dots in a column represents the community at a single timepoint. 76 strains are shown; they are colored according to their rank-order mean log_10_(relative abundance) across all samples in the experimental group (*n*=57). Both *Acidaminococcus* strains in the community are labeled; the removal of *Acidaminococcus* sp. D21 leads to an increase in *A. fermentans* DSM 20731, with most other strains staying at a similar level. (**D**) Removal of certain strains affected a large proportion of the community. The effect of a strain dropout on each strain was determined by calculating the *z*-score across all samples within an experimental group (Methods). The fraction of putative interactions was calculated based on the strains with |*z*|>2 for each strain dropout (*z*<-2 in blue, *z*>2 in red). Only strains above the limit of detection in the experimental group were counted, thus *n* is variable. (**E**) Some strains whose removal affected a large portion of the community were at high relative abundance, others low. The plot shows the mean log_10_(relative abundance) for each strain in the largest experimental group (*n*=57), excluding the sample in which that strain was dropped out. (**F**) A subset of the predicted interactions in the 104-member community can be recapitulated in binary culture. Interaction scores were high for some strain pairs (*a*→*b*, where removal of strain *a* affected strain *b*) with *z*<-2.6 (predicted positive interaction). We also included a pair with slightly smaller *z*-score (*z*=-2.1, *C. sporogenes* → *L. lactis*). Data are the mean of 2-6 replicates, and error bars represent 1 standard deviation from the mean. (**G**) *C. sporogenes* promotes *L. lactis* growth in binary culture. Growth curves are plotted for mono- or co-cultures of *C. sporogenes* and *L. lactis* in complete defined medium. Data are means and error bars represent 1 standard deviation from the mean (*n*=3). (**H**) Colony forming units for both strains in mono- or co-culture in complete defined medium. *C. sporogenes* levels were unaffected but the density of *L. lactis* increased ∼10-fold in co-culture (*p*-values correspond to Student’s t-tests (*n*=6). **: *p*<0.05.

We observed that the relative abundances in our data were correlated due to batch effects arising from the order in which communities were constructed (**Figure S6**); they clustered naturally into four sets. The two smallest sets (8 and 12 dropouts) were not considered further due to challenges in evaluating statistical significance. Within the two larger sets—60 and 24 dropouts—the relative abundances of each strain across samples were tightly distributed (**Figure 4B, S7**), enabling us to identify statistically significant responses using *z*-scores.

To test whether the single-strain dropout communities were deficient in the intended strain, we compared the relative abundance of each strain across all samples (**Figure 4B, Table S3**). For 71 of 84 communities, the strain we intended to remove had *z*<-3, indicating that the dropout was successful. In the remaining 13 communities, the intended strain was either below our limit of detection or was detected at 48 h; in both cases, the corresponding communities were not considered further (**Methods**).

Two additional filters were applied, resulting in the removal of six additional samples. First, five samples that had <10^6^ reads were not considered further due to the underrepresentation of low-abundance strains. Second, the *E. siraeum* dropout was missing an additional strain, *A. caccae*, and was removed from further consideration.

### Global analysis of strain-strain interactions

We analyzed the remaining 65 communities (**Table S4**) to identify strains whose relative abundance changed significantly in response to the absence of another strain. Putative interactions *a b* (where the arrow indicates that strain *b* increases or decreases in relative abundance in response to the absence of strain *a*) were considered further if they had |*z*|>2. We removed potentially spurious interactions that could have resulted from read mis-mapping or relative abundances near the lower limit of detection (**Methods**).

Despite the community’s complexity, large effects could be observed when certain strains were dropped out. For example, removing *Acidaminococcus sp*. D21 resulted in a 4-fold increase in the abundance of *Acidaminococcus fermentans* DSM 20731 (**Figure 4B**), presumably reflecting nutrient competition between strains of the same genus. The relative abundances of *A. fermentans* DSM 20731 were tightly distributed in other samples (**Figure 4A**, inset), so the change in relative abundance due to *Acidaminococcus sp*. D21 dropout was highly significant (*z*=5.1).

Most of the strain dropouts affected <5% of the other strains (**Figure 4C**). However, seven strains— when dropped out—impacted >10% of the remaining strains positively or negatively. Two of those strains, *Acidaminococcus sp*. D21 and *A. caccae* DSM 14662, were present at high relative abundance, so it is not surprising that relative abundances within the community redistribute upon their removal. However, the remaining strains—*Dorea formicigenerans* ATCC 27755, *Dialister invisus* DSM 15470, *Bacteroides dorei* 5_1_36/D4, *B. fragilis* 3_1_12, and *C. sporogenes* ATCC 15579—were present at a relative abundance of just ∼10^−3^–10^−2^, yet their removal altered the relative abundance of >7 other strains (**Figure 4D**). Conversely, four strains (*Bacteroides sp*. 2_1_16, *Bacteroides cellulosilyticus* DSM 14838, *Bacteroides ovatus* ATCC 8483, and *Mitsuokella multacida* DSM 20544) had relative abundances >1% but affected <2 strains when removed (**Figure 4D**), consistent with functional redundancy or another mechanism by which these strains are insulated from the rest of the community. Thus, the community is largely insensitive to strain removal even though a small subset of strains exhibit keystone-like properties.

### Validating candidate interactions in binary culture

Next, we tested whether interactions uncovered by the strain dropout screen could be observed in binary culture. Interactions can be direct or indirect (i.e. involve one or more additional strains), and context-independent or dependent (i.e. occur only in the background of a complex community). Only those interactions that are direct and context-independent would be reproducible in binary culture. We focused on interactions with *z*<-2 (rather than *z*>2), since the former involve growth promotion and are therefore simpler to validate in binary culture (**Table S4**).

We selected 32 candidate interactions with highly negative *z*-scores for further characterization. For these strain pairs, we compared the optical density of each strain when grown in monoculture versus co-culture, using the same defined medium as in the dropout screen. 23/32 strain pairs exhibited an increased carrying capacity relative to an additive growth model (**Figure 4E**) and/or decreased time to saturation in co-culture (**Figure S8**). These data indicate that the screen is an effective means of uncovering direct, context-independent strain-strain interactions. For 9/32 strain pairs, no interaction was observed in binary culture. While these negative results could be due to imperfections in the strain dropout experiment or analysis, they might also suggest that certain interactions can only be observed in the setting of a complex community.

We were intrigued that the level of *Lactococcus lactis* DSM 20729 decreased when *C. sporogenes* ATCC 15579 was dropped out of the community (*z*=-2.10). Although the effect size was small, the distribution of *L. lactis* relative abundances was particularly narrow, so the absence of *C. sporogenes* disrupted the growth of an otherwise context-insensitive strain.

To determine whether this interaction occurred in binary co-culture, we cultured *C. sporogenes* and *L. lactis* individually or together; *C. sporogenes* grew rapidly in defined medium while *L. lactis* was unable to grow on its own. The optical density of the co-culture was substantially higher than the sum of the optical densities of the individual cultures, suggesting that at least one of the strains grew more robustly in co-culture (**Figure 4F**). By counting colonies from the co-culture, we determined that *C. sporogenes* levels were unaffected but the density of *L. lactis* increased ∼10-fold (**Figure 4G**). These results validate the postulated interaction between *C. sporogenes* and *L. lactis* and suggest the possibility that the apparent sensitivity of *L. lactis* to leucine and arginine (**Figure 3E**) may in fact be a response to the drop in relative abundance of *C. sporogenes*. More broadly, our findings show that a growth stimulatory interaction between two strains can manifest even over one round of growth in the presence of 102 other strains.

## DISCUSSION

Our experimental system has three important features. First, by developing a community that is both defined and reasonably complex, we have generated a model system that will likely capture much of the biology of a native microbiome. Future refinements are needed, including additional bacterial strains to occupy unfilled niches as well as archaea, fungi, and viruses, all of which are important components of the native ecosystem.

Second, the computational pipeline we developed for read mapping makes it possible to analyze complex defined communities with high precision. Community structure can be quantified across six orders of magnitude in relative abundance, enabling the interrogation of low-abundance community members that play an important role in community function and dynamics (Buffie et al., 2015; Funabashi et al., 2020). The degree of technical and biological reproducibility (**Figure 2E**) is remarkable in a system this complex, which bodes well for future experimental efforts.

Third, the nutrient and strain dropout assays are a powerful format for probing the interactions that underlie community dynamics. The amino acid dropout screen tested 2,080 potential interactions (104 strains × 20 amino acids) using only 20 metagenomic sequencing samples, and the strain dropout screen tested 10,712 interactions (104 × 103 strains) using only 104 sequencing samples. Candidate interactions were observed in the background of a community and are therefore more likely to be relevant under native conditions; many of them were validated in co-culture experiments (**Figure 4E**). Taken together, our system for community assembly and measurement establish a framework for rapid, robust experimentation with complex consortia.

In its current form, our approach has two important limitations. First, when propagated *in vitro*, our community exhibits a different architecture than is observed *in vivo*. The consortium was dominated by Firmicutes that are typically found at lower relative abundances in the human gut, likely because the defined growth medium we used (SAAC) is rich in amino acids. Lowering the free amino acid content of the growth media and adding glycans, complex polypeptides, and host-derived factors such as bile acids could help steer the community toward a more typical *in vivo* architecture.

Second, the scale and complexity of our amino acid and strain dropout experiments precluded the possibility of carrying out multiple replicates. During the single passage over which communities were propagated, biomass increased by only 200-fold. As a result, the interactions we identified likely include some false positives; replicate experiments and additional growth passages could yield a larger or higher confidence set of interactions. Nonetheless, the strain-amino acid and strain-strain interactions we validated show that the experiments were sufficiently sensitive and reproducible to uncover real interactions.

The properties of this community in the context of host colonization are described in an accompanying manuscript. Together, these efforts constitute a starting point for a defined, full-scale model system for the gut microbiome. Such a system would yield great dividends, as the research communities around yeast, worms, flies, and mice have shown over decades. It would be even more powerful in conjunction with new genetic tools to manipulate specific members of this community.

## STAR★METHODS

Detailed Methods are provided in the online version of this paper and include the following:

- KEY RESOURCES TABLE
- RESOURCE AVAILABILITY
  ○ Lead contact
  ○ Materials availability
  ○ Data and code availability
- EXPERIMENTAL MODEL AND SUBJECT DETAILS
  ○ Bacterial strains and culture conditions
- METHOD DETAILS
  ○ Metagenomic sequencing
  ○ Metagenomic read mapping
  ○ Amino-acid dropout experiments
  ○ Creation of *C. sporogenes* mutants
  ○ ATP assay
  ○ Strain-dropout experiments
  ○ Pairwise co-culture growth measurements
- QUANTIFICATION AND STATISTICAL ANALYSIS

## STAR★METHODS

### Lead contact

Further information and requests for resources and reagents should be directed to and will be fulfilled by the lead contact, Michael Fischbach.

### Materials availability

*C. sporogenes* mutants are available on request. The strains used in this study are available from the sources listed in the Key Resource Table.

### Data and code availability

Metagenomic and whole-genome sequencing datasets generated for this study will be available at the Sequence Read Archive at the time of publication. Ninjamap is available at https://github.com/FischbachLab/ninjamap and the associated docker containers are available at https://hub.docker.com/orgs/fischbachlab/repositories.

## EXPERIMENTAL MODEL AND SUBJECT DETAILS

### Bacterial strains and culture conditions

Bacterial strains were selected based on metagenomic sequencing data from the NIH Human Microbiome Project (Kraal et al., 2014). The mean relative abundance and prevalence of each strain were quantified using the 81 samples from healthy human patients from North America. The ∼200 strains that appeared in ≥37 of the 81 samples were considered for inclusion in the community. We were able to obtain 104 of these strains from public repositories (Key Resources Table).

Strains were cultured in anaerobic conditions (10% CO_2_, 5% H_2_, 85% N_2_) in 2 mL 96-well plates for 24-48 h in their respective growth media (**Table S1**): Mega Medium (McNulty, et. al. 2015) supplemented with 400 µM Vitamin K2, or Chopped Meat Medium supplemented with Mega Medium carbohydrate mix (McNulty, et. al. 2015) and 400 µM Vitamin K2. For strain storage, 200 µL of liquid culture was aliquoted 1:1 into sterile 50% glycerol in a 1 mL 96-well plate. The plate was covered with an airtight silicone fitted plate mat, edges were sealed with O_2_-impervious yellow vinyl tape, and the plate was frozen at -80 °C. To revive cultures, the storage plate was defrosted in the anaerobic chamber and 100 µL from each well was used to inoculate 900 µL of appropriate fresh medium. Twenty-four hours post-revival, each well was visually inspected and wells that did not exhibit obvious growth were re-inoculated with an additional 100 µL of frozen stock. Each storage plate included 3-4 “sentinel” wells containing only growth medium that were used to monitor potential contamination.

## METHOD DETAILS

### Constructing high quality genome assemblies

We obtained the latest RefSeq (O’Leary et al., 2016) assembly for each strain in our community and assessed its quality based on contig statistics from Quast (Gurevich et al., 2013) v. 5.0.2 and SeqKit (Shen et al., 2016) v. 0.12.0, using GTDB-tk (Chaumeil et al., 2019) v. 1.2.0 for taxonomic classification. A linear combination of the completeness and contamination scores (completeness–5×contamination) derived from the CheckM (Parks et al., 2015) v. 1.1.2 lineage workflow was used along with the other metrics to include or exclude genomes in the GTDB (Parks et al., 2018, 2020) release 89 database (https://gtdb.ecogenomic.org/faq#gtdb_selection_criteria). Genomes that contained any number of Ns, contained over 100 contigs, contained GTDB lineage warnings or multiple matches, or had CheckM completeness <90, contamination >10, and combination score <90 were resequenced and reassembled. Our hybrid assembly pipeline contains a workflow for *de novo* and reference-guided genome assembly using both Illumina short reads and PacBio or Nanopore long reads. The workflow has three main steps: read pre-processing, hybrid assembly, and contig post-processing. Read pre-processing included 1) quality trimming/filtering (bbduk.sh adapterFile=“adapters,phix” k=23, hdist=1, qtrim=rl, ktrim=r, entropy=0.5, entropywindow=50, entropyk=5, trimq=25, minlen=50), with adaptors and phix removed with kmer right trimming, kmer size of 23, Hamming distance 1 (allowing one mismatch), quality trimming of both sides of the read, filtering of reads with an average entropy <0.5 with entropy kmer length of 5 and a sliding window of 50, trimming to a Q25 quality score, and removal of reads with length <50 bp; 2) deduplication (bbdupe.sh); 3) coverage normalization (bbnorm.sh min=3) such that depth <3x was discarded; 4) error correction (tadpole.sh mode=correct); and 5) sampling (reformt.sh). All pre-processing was carried out using BBtools v. 38.37 for short reads. For long reads, we used filtlong v. 0.2.0 (fitlong --min_length 1000 --keep_percent 90 --length_weight 10) to discard any read <1 kb and the worst 10% of read bases, as well as to weigh read length as more important when choosing the best reads. Hybrid assembly was performed by Unicycler (Wick et al., 2017) v. 0.4.8 with default parameters using pre-processed reads. After assembly, the contigs from the assembler were scaffolded by LRScaf (Qin et al., 2018) v. 1.1.9 with default parameters. If the initial assembly did not produce the complete genome, gaps were filled by long reads TGS-GapCloser (Xu et al., 2019) v. 1.0.1 with default parameters.

If no long reads were available, short paired-end reads were assembled *de novo* using SPAdes (Bankevich et al., 2012) v. 3.13.1 with the --careful option to reduce the number of mismatches and short indels during assembly of small genomes. Assembly quality was assessed based on the CheckM v. 1.1.2 lineage. If contamination was detected, contigs corresponding to the genome of interest were extracted from the contaminated assembly using MetaBAT2 (Kang et al., 2019) v. 2.2.14 with default parameters.

Finally, the assembled genomes were evaluated using the same criteria as the RefSeq assemblies, and the assembly for each species with the best overall quality metrics was chosen as the reference assembly. This procedure resulted in the replacement of eight genomes: two from a PacBio/Illumina hybrid assembly, one from a Nanopore/Illumina hybrid assembly, one from a reference-guided Illumina assembly, and four from short-read assemblies of the respective isolate samples followed by binning (**Table S5**).

### Generating simulated sequencing reads

*In silico* data were generated to evaluate the Ninjamap algorithm in the absence of genome assembly errors and sequencing quality issues. Grinder (Angly et al., 2012) v. 0.5.4 was applied to each genome to generate error-free reads with the following parameters: -read_distribution 140, -insert_size 800, -mate_orientation FR, -delete_chars ‘-∼*NX’, -mutation_dist uniform 0, -random_seed 1712, - abundance_model uniform, -qual_levels 33 31, -fastq_output 1. The -coverage_fold parameter was adjusted based on the cases described below.

#### Uniform abundance isolate dataset

This dataset was created to test the sensitivity and specificity of the algorithm against our database of genomes. *In silico* data were generated for each genome with uniform coverage of 10X or 100X. We were able to consistently identify the correct genome regardless of coverage (**Figure 2C**). Some cross mapping was observed at ∼0.01% relative abundance, likely because some genomes in our database shared more than 99% average nucleotide identity, making cross-mapping unavoidable. Thus, we generally treat 10^−4^ as a conservative lower bound for confident relative abundance estimation.

#### Variable abundance community dataset

*In silico* reads were generated for each genome at 10X, 0.1X, and 0.001X uniform coverage. Three datasets of mixed community reads were generated including every genome at a coverage randomly selected from the three levels. The observed relative abundance of each genome in our database was calculated using the NinjaMap algorithm and compared to the expected relative abundance based on coverage level, which ranged from ∼3×10^−6^ to 0.03. We could estimate relative abundances accurately for genomes present at >10^−5^ (**Figure 2D**). However, for lower relative abundances, we observed some discrepancies corresponding to the same genomes with mis-mapping against isolate datasets, indicating that high similarity between genomes begins to confound the algorithm at very low relative abundances.

### Metagenomic sequencing

The same experimental pipeline was used for sequencing bacterial isolates and synthetic communities. Bacterial cells were pelleted by centrifugation under anaerobic conditions. Genomic DNA was extracted using the DNeasy PowerSoil HTP kit (Qiagen) and quantified in 384-well format using the Quant-iT PicoGreen dsDNA Assay Kit (Thermofisher). Sequencing libraries were generated in 384-well format using a custom low-volume protocol based on the Nextera XT process (Illumina). Briefly, the concentration of DNA from each sample was normalized to 0.18 ng/µL using a Mantis liquid handler (Formulatrix). If the concentration was <0.18 ng/µL, the sample was not diluted further. Tagmentation, neutralization, and PCR steps of the Nextera XT process were performed on a Mosquito HTS liquid handler (TTP Labtech), leading to a final volume of 4 µL per library. During the PCR amplification step, custom 12-bp dual unique indices were introduced to eliminate barcode switching, a phenomenon that occurs on Illumina sequencing platforms with patterned flow cells (Sinha et al., 2017). Libraries were pooled at the desired relative molar ratios and cleaned up using Ampure XP beads (Beckman) to achieve buffer removal and library size selection. The cleanup process was used to remove fragments <300 bp or >1.5 kbp. Final library pools were quality-checked for size distribution and concentration using a Fragment Analyzer (Agilent) and qPCR (BioRad). Sequencing reads were generated using a NovaSeq S4 flow cell or a NextSeq High Output kit, in 2×150 bp configuration. 5-10 million paired-end reads were targeted for isolates and 20-30 million paired-end reads for communities.

### Generating and normalizing the NinjaMap database

The first step in the pipeline was to assess the uniqueness of each genome in the community. We generated error-free *in silico* reads such that each genome was uniformly covered at 10x depth. Each such genome read set was aligned to all genomes in the community. The uniqueness of a genome was defined as the fraction of the genome that did not have reads cross-mapped from another strain; uniqueness values were between 0 and 1, such that more unique genomes have a value closer to 1. The uniqueness value of a strain was used to normalize its final relative abundance in any community sample. All genome sequences were combined into one fasta file and a Bowtie2 (Langmead and Salzberg, 2012) v. 2.3.5.1 index was computed for future alignments. The database and strain weights were recomputed each time the community or a genome was updated.

### Metagenomic read mapping

Paired-end reads from each sample were aligned to the database using Bowtie2 with maximum insert length (-maxins) 3000, maximum alignments (-k) as 300, suppressed unpaired alignments (--no-mixed), suppressed discordant alignments (--no-discordant), suppressed output for unaligned reads (--no-unal), required global alignment (--end-to-end), and using the “--very-sensitive” alignment preset. The output was processed in Samtools (Li et al., 2009) to convert the alignment output from SAM output stream to BAM format. The BAM file was sorted and indexed by coordinates.

### NinjaMap alignment scoring

A primary goal of the NinjaMap algorithm is to analyze and tabulate every input read. A successful match was defined as a read aligned to a genome at 100% identity across 100% of the read length. If a read was uniquely matched to a single strain, its mate pair was also recruited as long as it had at least one match to the same strain. If exactly 1 strain was a perfect match for both reads, the pair was considered a “primary pair” and a score of 1 was given for each read. If >1 or 0 strains were a match for both reads, both reads were placed in escrow and analyzed separately as described below.

By prioritizing paired-read scoring, noise was significantly reduced while ensuring that as many reads as possible were considered for abundance estimates. Once preliminary strain abundances were calculated based on primary pairs, reads in escrow were then assigned fractionally to the strains to which they aligned perfectly. The fractional assignment was calculated based on the primary read abundances of each strain, normalized by the size of the unique region of each genome within the database, such that the total contribution for a read was 1. In some cases, an individual escrowed read matched to a strain without any matches to primary pairs; such reads were discarded and not used in the final estimates.

Finally, the total score for each strain in the database was normalized by the number of reads that aligned to the database, so that the relative abundances of all strains summed to 1.

### Amino acid dropout experiments

Strains were passaged by diluting 1:10 into fresh growth medium every 24 h for 2-3 days. The day before amino acid dropout experiments, cultures were diluted 1:10 into 1 mL of fresh medium and grown for 24 h as inoculation working stocks. To measure OD_600_, strains were diluted 1:10 into 150 µL of the appropriate culture medium and a plate reader was used to measure absorbance at 600 nm. Stocks were diluted to a final OD_600_ of 0.1 using fresh growth medium. If a culture did not reach OD_600_ of 0.1, the entire culture was used as the working stock for community assembly. Equal volumes of each stock were pooled to create a 104-member synthetic community. The community was centrifuged at 5000 x *g* for 5 min, washed, and resuspended in an equivalent volume of PBS to generate the pooled community working stock. SAAC medium (Dodd et al., 2017) was made containing all amino acids at 1 mM concentration except for cysteine, which was added at 4.126 mM (**Table S6**). Twenty similar media were made in which one amino acid at a time was removed. 1.6 mL of each medium was aliquoted in triplicate and inoculated with the pooled community at 1:10 or 1:100 dilution. Four 100 µL aliquots of each culture were collected at 48 h and processed for metagenomic sequencing.

### Constructing *C. sporogenes* mutants

*C. sporogenes* deletion mutants were constructed using a previously reported protocol (Guo et al., 2019); the strains and primers used for each mutant are listed in **Table S7**. In brief, from plasmids CS_OTC and CS_ADI—which harbor the targeting and repair templates—we amplified DNA sequences encoding the gRNA locus (the gRNA plus adjacent elements and the repair template) and ligated the amplicon into the pMTL82254 backbone. These repair templates consist of 700- to 1200-bp sequences flanking the 40- to 100-bp sequence targeted for excision.

To construct the Δ*adi* strain, a gRNA fragment was purchased from Quintara and amplified with primers fwd_pMTL82254_NotI, rev_gRNA_flank1. The two flanking regions were amplified from *C. sporogenes* genomic DNA using the primers 5rev_flank1 and 5fwd_flank1_flank2 for flank 1 and 5rev_flank1_flank2 and 5fwd_flank1_flank2 for flank 2. Next, the flanking regions were joined by amplifying with primers fwd_gRNA_flank1 and rev_flank2. The amplified gRNA fragment was attached to the joined flank construct by amplifying with primers fwd_pMTL82254_NotI and rev_pMTL82254_AscI. Finally, the pMTL82254 plasmid and the construct containing the gRNA, flank1, and flank2 regions were digested with NotI and AscI and ligated with T4 ligase (NEB). The final construct was named CS_ADI.

To make the Δ*otc* strain, the gRNA fragment was purchased from Quintara and amplified with fwd_pMTL82254_NotI and rev_OTC_gRNA_flank1. The two flanking regions were amplified from *C. sporogenes* genomic DNA using the primers fwd_OTC_gRNA_flank1 and rev_OTC_flank1_flank2 for flank 1 and fwd_OTC_flank1_flank2 and rev_OTC_flank2 for flank 2. Next, the flanking regions were joined by amplifying with the primers fwd_OTC_gRNA_flank1 and rev_OTC_flank2. The amplified gRNA fragment was attached to the joined flank construct by amplifying with fwd_pMTL82254_NotI and rev_pMTL82254_AscI. Finally, the pMTL82254 plasmid and the construct containing the gRNA, flank1, and flank2 regions were digested with NotI and AscI and ligated with T4 ligase (NEB). The final construct was named CS_OTC.

CS_OTC or CS_ADI was electroporated into *Escherichia coli* S17 cells and conjugated into *C. sporogenes* strain ATCC 15579 using a previously described method (Guo et al., 2019). In brief, a single colony of wild-type *C. sporogenes* was used to inoculate 2 mL of TYG broth (3% (w/v) tryptone, 2% (w/v) yeast extract, 0.1% (w/v) sodium thioglycolate) and incubated anaerobically in an atmosphere consisting of 10% CO_2_, 5% H_2_, and 85% N_2_. *E. coli* S17 cells with CS_OTC or CS_ADI were grown in LB broth supplemented with 250 μg/mL erythromycin at 30 °C with shaking at 225 rpm. After 17-24 h, 1 mL of this culture was centrifuged at 1000 x *g* for 1 min and washed twice with 500 µL of PBS (40 mM potassium phosphate, 10 mM magnesium sulfate, pH 7.2). The pellet was transferred into the anaerobic chamber and 250 µL of *C. sporogenes* overnight culture was added and mixed with the cell pellet. 30 µL aliquots of the mixture were plated on a pre-reduced TYG agar plate in eight spots. The plate was tilted to coalesce the spots and incubated for 24 h. Biomass from the plate was scraped using a sterile inoculation loop and suspended in 250 µL of pre-reduced PBS. 100 µL of the cell suspension was plated on TYG agar containing 10 µg/mL erythromycin and 250 μg/mL D-cycloserine to isolate single colonies. One colony was picked, sequence verified, and used as the starting point for the next conjugation.

In the second conjugation, *E. coli* S17 cells containing pMTL83153_fdx_Cas9 were grown in LB broth supplemented with 25 μg/mL chloramphenicol at 30 °C with shaking at 225 rpm. After washing, the pellet was moved into the anaerobic chamber and 250 μL of an overnight culture of *C. sporogenes* harboring the CS_OTC vector was thoroughly mixed with the *E. coli* cell pellet. 30 µL aliquots of the mixture was plated on a pre-reduced TYG agar plate in eight spots. The plate was tilted to coalesce the spots and incubated for 72 h. Biomass from the plate was scraped using a sterile inoculation loop and resuspended in 250 μL of pre-reduced PBS. 100 µL of the cell suspension was plated on each of two pre-reduced TYG agar plates containing 10 μg/mL erythromycin, 15 μg/mL thiamphenicol, and 250 μg/mL D-cycloserine. *C. sporogenes* colonies typically appeared after 36-48 h, and 8-10 colonies were re-streaked on pre-reduced TYG agar plates containing 10 μg/mL erythromycin, 15 μg/mL thiamphenicol, and 250 μg/mL D-cycloserine to isolate single colonies. The isolated colonies were used to inoculate pre-reduced TYG broth supplemented with 10 μg/mL erythromycin and 15 μg/mL thiamphenicol, and genomic DNA was isolated using a Quick DNA fungal/bacterial kit (Zymo Research). Primers ADI_532_fwd and ADI_22_rev or OTC_5_up_fwd and OTC_930_down_rev (**Table S2**) were used to verify deletions.

### ATP assay

An aliquot from a frozen stock of *C. sporogenes* was used to inoculate 5 mL of TYG broth and grown to stationary phase (∼24 h). Cells were diluted 1:1000 into 20 mL of TYG broth and grown to late-log phase (∼16 h). Cells were harvested by centrifugation (5,000 x *g* for 10 min at 4 °C) and washed twice with 20 mL of pre-reduced PBS. 100 µL of cells was seeded into rows of a 96-well microtiter plate (12 wells per condition). 200 µL of pre-reduced 2 mM substrate (arginine) in phosphate washing buffer, or 200 µL of buffer alone, were dispensed into rows of a separate 96-well microplate. At *t*=0, 100 µL of substrate or buffer were added to the cells and mixed gently by pipetting. At *t*=-5 min, -1 min, 30 s, 1 min, 2 min, 5 min, 10 min, 20 min, 30 min, 45 min, 60 min, and 90 min, 10 µL of cells were extracted and mixed with 90 µL of DMSO to quench the reaction and liberate cellular ATP. For the time points *t*=-5 min and -1 min—prior to the addition of buffer or substrate—5 µL of cell suspension was harvested and 5 µL of either buffer or substrate were added to the cell-DMSO mixture to bring the total volume to 100 µL. The ATP content from 10 µL aliquots of lysed cells was measured using a luminescence-based ATP determination kit (Invitrogen, Cat. #A22066). Absolute ATP levels were calculated using a calibration curve with known concentrations of ATP.

### Strain dropout experiments

Strains were passaged by diluting 1:10 into fresh growth medium every 24 h for 2-3 days. The day before strain dropout experiments, cultures were diluted 1:10 into 1 mL of medium and grown overnight as inoculation working stocks. To measure OD_600_, strains were diluted 1:10 into 150 µL of the appropriate culture medium and a plate reader was used to measure absorbance at 600 nm. Stocks were normalized using fresh growth medium to a final OD_600_ of 0.1. If an overnight culture did not reach OD_600_ of 0.1, the entire culture was used as the working stock for community assembly. For community assembly, 10 mL of each stock were pipetted into a 12-well reservoir plate.

Strain dropouts were performed on a row-by-row basis in a 96-well deep-well plate. For example, to individually dropout each of the 12 strains in row A, equal volumes of all community members not in row A were pooled and aliquoted into 12 sterile 1.5 mL Eppendorf tubes. This process resulted in 120 µL of each of 104-12=92 strains for each of 12 communities being combined into a 11.04 mL pool, which was divided into 12 aliquots of 920 µL in sterile Eppendorf tubes. To add the strains in row A, 12 sub-communities were created by pooling 10 µL of each of the 12 strains except for the intended dropout. 10 µL of PBS were added, for a total volume of 120 µL, which was added to the 920 µL pooled community lacking row A, resulting in the creation of 12 communities with volume 1040 µL, each lacking one of the 104 strains. This process was repeated for all rows to cover the 104 strains over 2 days; 24 and 80 dropouts were constructed on day 1 and 2, respectively. Each dropout community was washed once via centrifugation at 5000 x *g* for 5 min and resuspended in an equal volume of sterile PBS. Sixteen microliters of each dropout community were used to inoculate 1.6 mL of SAAC medium. Four hundred microliter aliquots were collected at 12 h, 24 h, and 48 h post-inoculation. The initial dropout community stocks and all aliquots were processed for metagenomic sequencing using a Power Soil DNA extraction kit (Qiagen).

### Pairwise co-culture growth measurements

Each strain was inoculated from a frozen stock into its optimal growth medium (Mega Medium or Chopped Meat Medium) and the culture was incubated for 24 h at 37 °C. OD_600_ was measured, and 1 mL of each strain was washed with sterile 1x PBS. 0.75 µL of each strain (for co-cultures) or PBS (for monocultures) was added to 148.5 µL SAAC medium for a total volume of 150 µL. The baseline OD_600_ and growth curves were measured using an Epoch plate reader (Biotek). For *C. sporogenes* and *L. lactis*, cultures were normalized to OD_600_=1.0 before mixing. All cultures were performed in technical triplicate with 2-6 biological replicates. For each growth curve, the interaction score α was computed based on an additive null model (Aranda-Díaz et al., 2020):

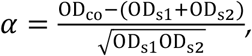

where OD_co_, OD_s1_, and OD_s2_ are the maximum OD values of the co-culture and the individual strains s1 and s2, respectively. The time to reach half maximum absorbance (*t*_1/2_) was measured for individual cultures and co-cultures and labeled as significant if average co-culture *t*_1/2_ was greater than either individual average *t*_1/2_. Student’s t-test was utilized to determine significant changes with *p*<0.05.

### Mis-mapping estimation using monoculture sequencing

Read fractions were analyzed using custom Matlab (Mathworks, R2018a) code. Read fractions were rescaled to sum to 1, thereby reflecting the relative abundances of reads mapped to one of the 104 genomes in our database. These data were used to calculate expected relative abundances of each strain from mis-mapping data as described below. The expected mis-mapped relative abundance (*E*_*i,j*_) for strain *i* in a sample *j* was calculated as

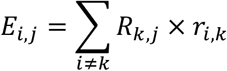

where *R*_*k,j*_ is the relative abundance of strain *k* in sample *j* and *r*_*i,k*_ is the relative abundance of strain *i* in sequencing data of strain *k* as an isolate. Strains whose relative abundance was comparable to the expected mis-mapped relative abundance (*E*_*i,j*_<10*r*_*i,j*_) were removed from further analysis.

### Amino acid dropout data analysis

Read fractions were rescaled to sum to 1, thereby reflecting the relative abundances of reads mapped to one of the 104 genomes in our database. The effect of removal of an amino acid on a strain was estimated by calculating the z score 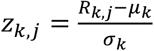, where *R*_*k,j*_ is the log_10_(relative abundance) of strain *k* in sample *j* and *μ*_*k*_ and *σ*_*k*_ are the mean and standard deviation, respectively, of log_10_(relative abundance) for strain *k* across all samples except the cysteine dropout. The cysteine dropout sample was excluded from the calculation of *µ*_*k*_ and *s*_*k*_ because this sample was an obvious outlier. Data points that could be explained by mismapping were removed. Putative interactions were identified based on |*z*_*j,k*_|>2, i.e. amino acid dropouts that changed the log_10_(relative abundance) of strain *k* by ≥2 standard deviations relative to its mean. Some strains varied in relative abundance by several orders of magnitude; as a result, *σ*_*k*_ was large, so putative interactions would be missed using *z*-scores.

To identify clusters of strains that responded similarly or amino acids that elicited a similar response, we normalized *R*_*k,j*_ for each strain across samples by subtracting *µ*_*k*_ and performed hierarchical clustering of both strains and amino acid dropouts on a dataset including strains that were detected in all 21 amino acid dropout samples.

### Strain interactions in strain-dropout data

Read fractions were rescaled to sum to 1, reflecting the relative abundances of reads mapped to one of the 104 genomes in our database. Samples with <10^6^ reads, as well as the *Eubacterium siraeum* DSM 15702 dropout (which appeared to be missing an additional strain, *A. caccae*), were discarded from further analysis.

To identify batch effects that caused unintentional groupings within the dataset, we subtracted the mean log_10_(relative abundance) of each strain in *t*=48 h samples from the log_10_(relative abundance) in each sample. These normalized relative abundances were used to calculate the Pearson correlation coefficient of each pair of samples. Based on this correlation matrix, samples were split into 4 experimental groups that corresponded to natural groupings arising from the experimental setup (**Figure S6**), with group sizes *n*=21, 57, 12 and 8. For the two largest groups, *z*-scores were calculated for each strain based on the mean and standard deviation of log_10_(relative abundances) within the corresponding experimental group. To account for noise in low-abundance strains, *z*-scores were only calculated for strains that were undetected in <5 samples in the experimental group. Data points that could be explained by mismapping were removed. Statistics were calculated using only detected strains in each sample; undetected strains were only set to an arbitrary small number for graphical representation. Putative interactions were defined based on |*z*_*j,k*_|>2.

### Statistical analysis

The statistical details of experiments can be found in the figure legends. Reported *n* values are the total samples (cultures) per group. Unless otherwise stated, *p*-values were not corrected for multiple hypothesis testing. Benjamini-Hochberg corrections, hypergeometric tests, Student’s t-tests (unpaired or two-tailed), and Kruskal-Wallis tests were performed in MATLAB.

## ACKNOWLEDGMENTS

We are deeply indebted to members of the Fischbach and Huang labs for helpful discussions, and to Rod Mackie (UIUC) for bacterial strains used in this study. A.A.-D. is a Howard Hughes Medical Institute International Student Research fellow, a Stanford Bio-X Bowes fellow, and a Siebel Scholar. This work was supported by the Stanford Microbiome Therapies Initiative (M.A.F., K.C.H.), NIH grants RM1 GM135102 (K.C.H.), DP1 DK113598 (M.A.F.), P01 HL147823 (M.A.F.), and R01 DK101674 (M.A.F.), the Bill and Melinda Gates Foundation (M.A.F.), an HHMI-Simons Faculty Scholars Award (M.A.F.), the Leducq Foundation (M.A.F.), the Stanford-Coulter Translational Research Grants Program (M.A.F.), the Chan Zuckerberg Biohub (K.C.H., M.A.F.), and the Allen Discovery Center at Stanford on Systems Modeling of Infection (K.C.H.).

## AUTHOR CONTRIBUTIONS

Conceptualization: A.C., M.A.F. Methodology and investigation: A.C., A.A.-D., S.J., F.Y., M.I., X.M., A.P., A.W., A.L.S., A.D., K.C.H., M.A.F. Formal analysis: A.C., A.A.-D., S.J., F.Y., M.I., X.M., K.C.H., M.A.F. Visualization: A.C., A.A.-D., S.J., K.C.H., M.A.F. Supervision: K.C.H., M.A.F. Writing: A.C., A.A.-D., S.J., F.Y., K.C.H., M.A.F. All authors reviewed the manuscript before submission.

