## Supplementary Information for "Systematic dissection of a complex gut bacterial community"

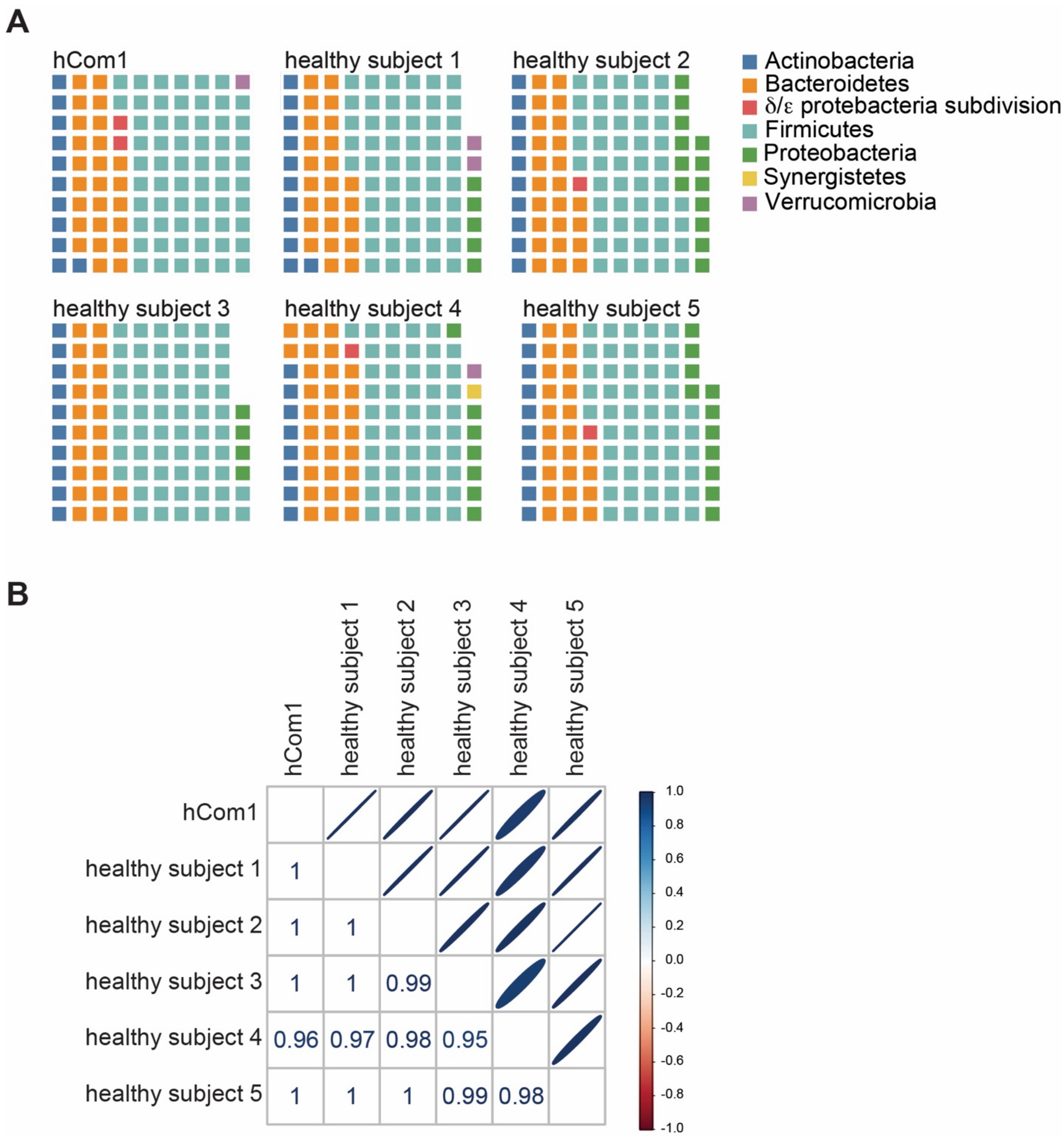


**Figure S1: The 104-member community resembles the phylogenetic distribution of a typical Western human gut community.** (**A**) A waffle plot comparing the phylogenetic distribution of strains from our synthetic community (hCom1) versus fecal communities from five healthy human donors. The number of squares represents the percentage of strains in each phylum as computed by MIDAS. The number of squares in some grids is <100 because some reads do not map to a known phylum. Phyla with <0.5% of the total reads are not represented.

(**B**) Correlation coefficients between the synthetic community and each healthy human fecal community.

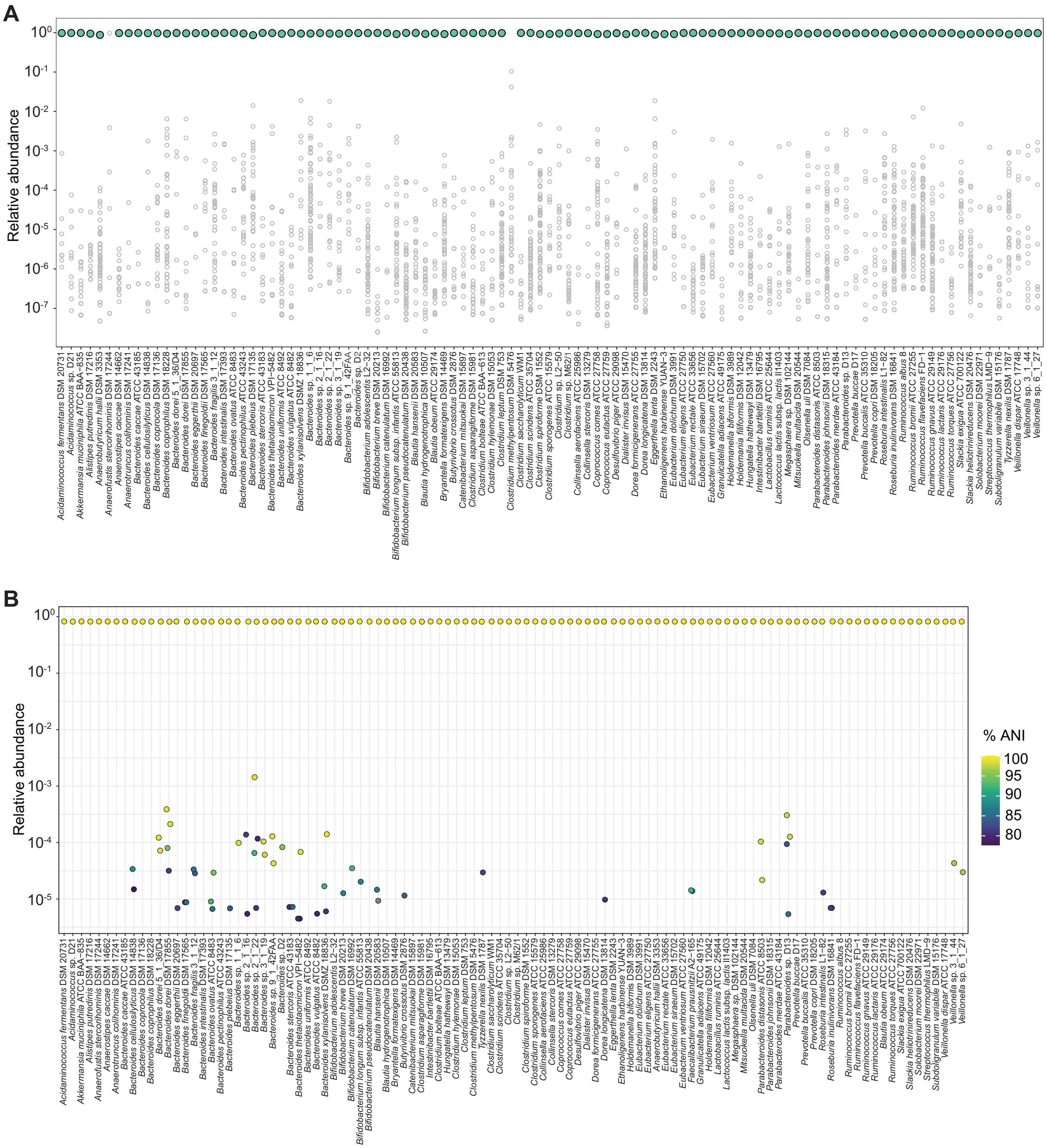


**Figure S2: Performance of NinjaMap on simulated and actual single-strain sequencing data.** (**A**) NinjaMap analysis of data derived from sequencing each strain in the community individually. The green dot represents the relative abundance of the strain that was sequenced; gray dots represent the relative abundance of other strains in the sample. In general, the rate of off-mapping was below $10^{-3}$. (**B**) NinjaMap analysis of simulated (*in silico* generated) sequencing data for each strain in hCom1. The yellow dot represents the relative abundance of the strain that was sequenced; the remaining dots represent the relative abundance of other strains that were called in the sample. Points are colored based on % average nucleotide identity (ANI) values compared to the strain being sampled, and a small amount of jitter has been added to aid in viewing overlapping circles. Mapping to the correct genome was achieved for the vast majority of strains, even when other highly similar genomes are present in the database.

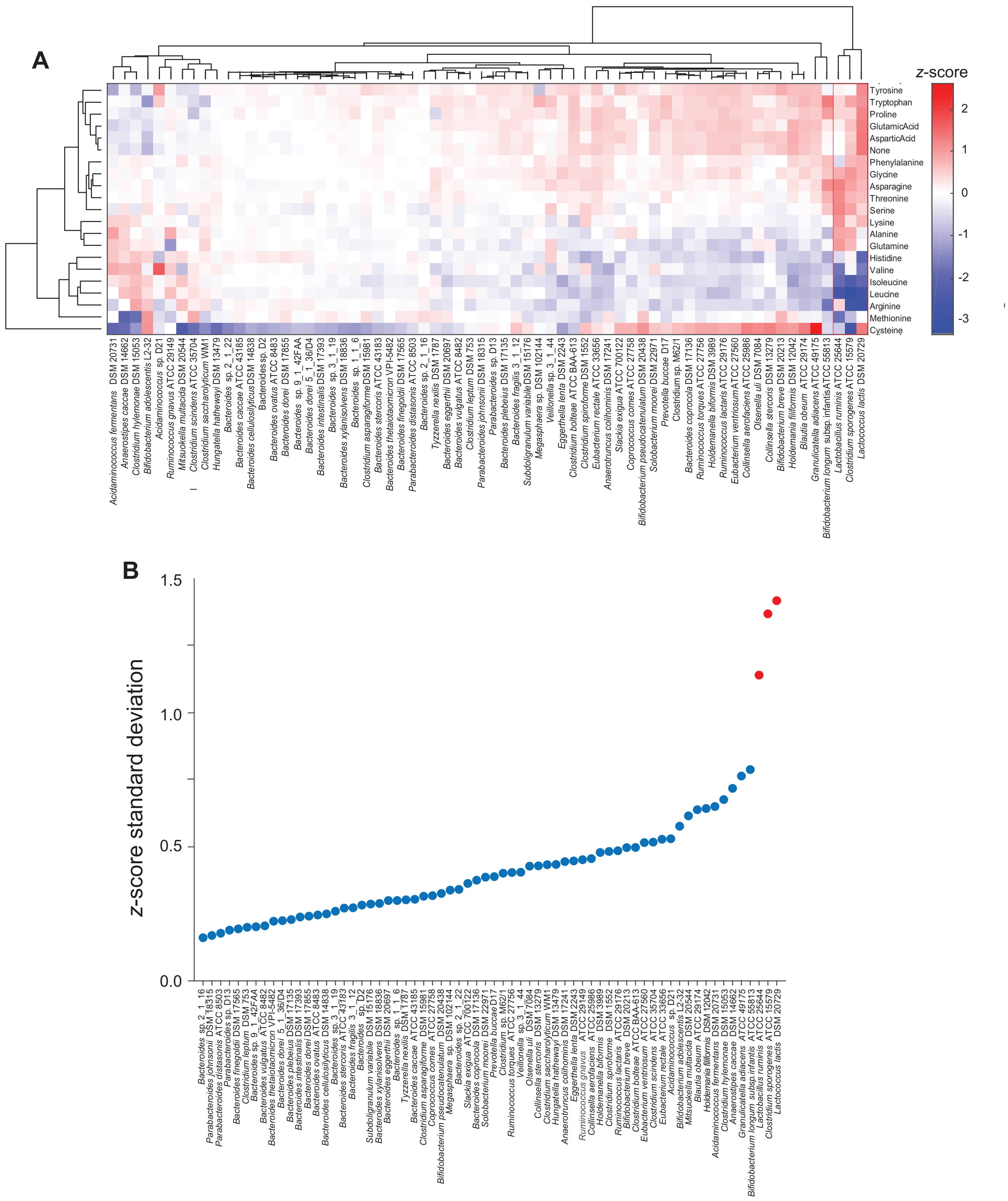


**Figure S3: *C. sporogenes*, *L. lactis*, and *L. ruminis* are highly sensitive to amino acid perturbation.** (**A**) Hierarchical clustering of strains (*x*-axis) and samples (amino acid dropouts, *y*-axis) revealed that *C. sporogenes*, *L. lactis*, and *L. ruminis* were sensitive to the removal of branched chain amino acids. The *z*-score in each condition is plotted. (**B**) The standard deviation of the *z*-scores of *C. sporogenes*, *L. lactis*, and *L. ruminis* (red) was substantially higher than of all other strains, indicating that they are more highly variable across amino acid dropouts.

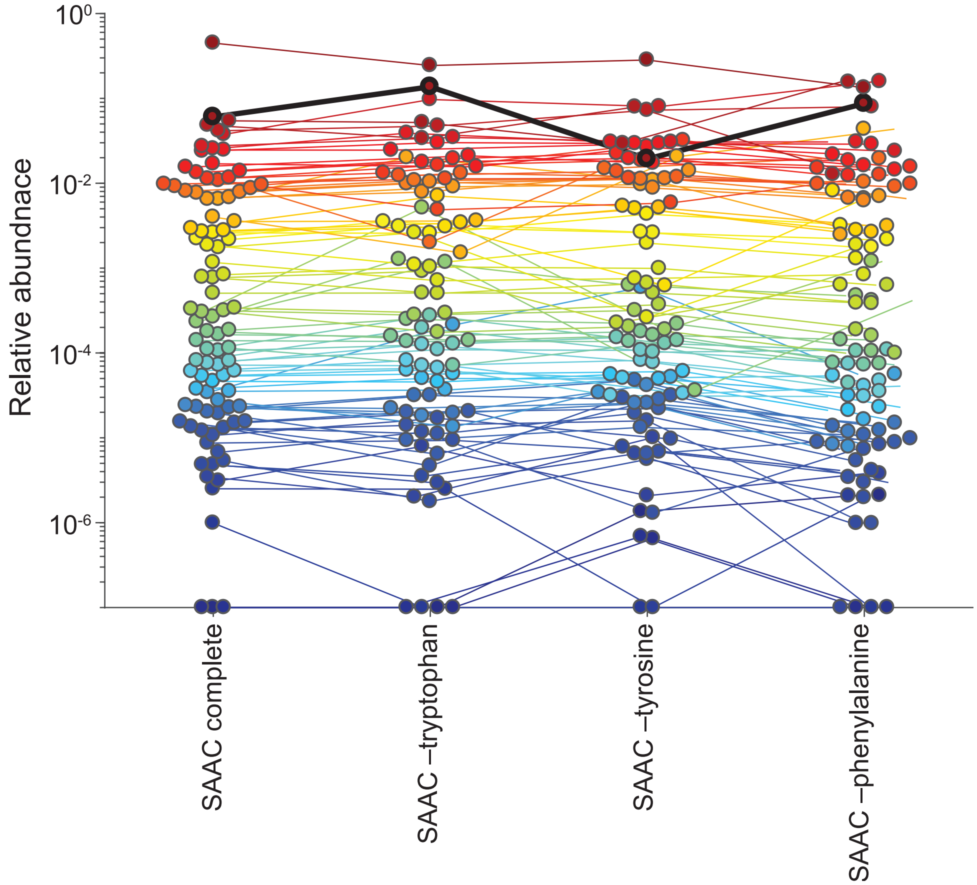


**Figure S4: The relative abundance of *C. sporogenes* was not affected significantly by the removal of phenylalanine, tyrosine, or tryptophan.** Each dot is an individual strain; the collection of dots in a column represents the community at 48 h. Strains are colored according to their rank-order abundance in the community grown in complete defined medium. Undetected strains were set to $10^{-7}$ for visualization. The thick black line shows the relative abundance of *C. sporogenes*.

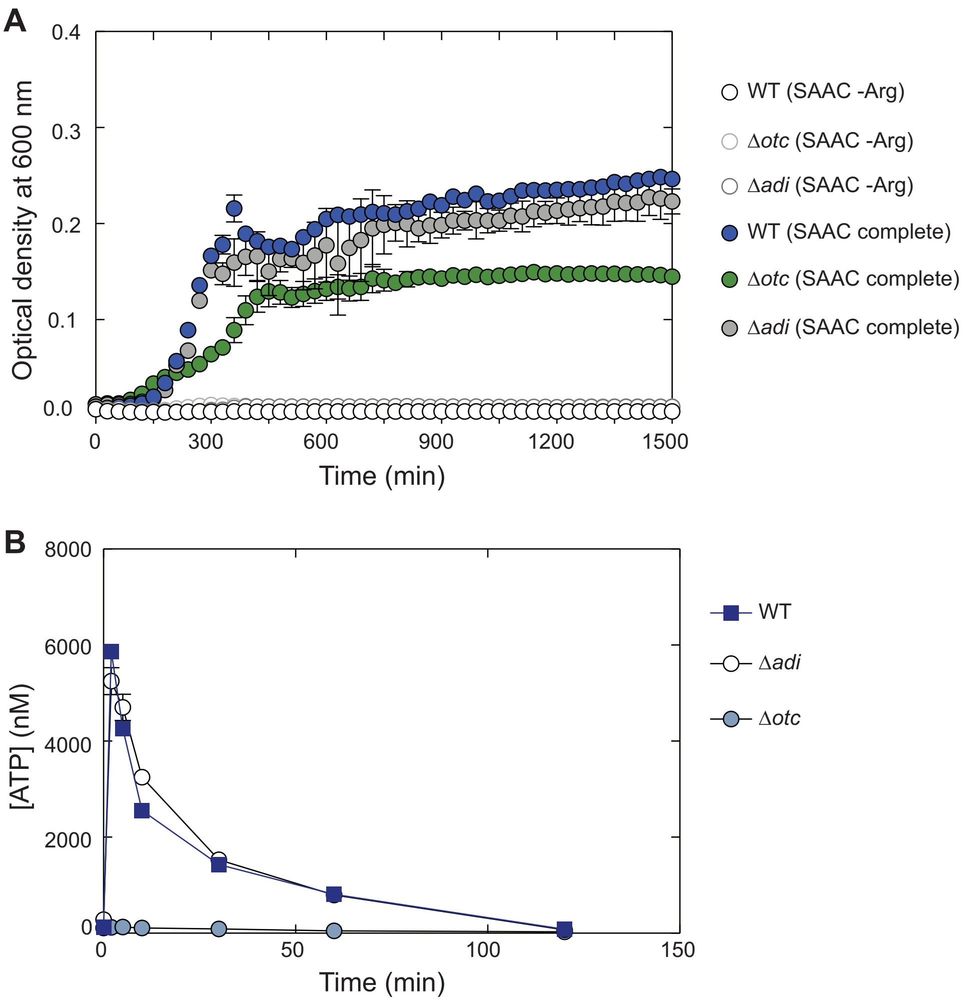


**Figure S5: Deletion of ornithine transcarbamoylase (*otc*), but not arginine deiminase (*adi*), impairs arginine-dependent growth in minimal medium and ATP production.** (**A**) Wild-type *C. sporogenes* and Δ*adi* and Δ*otc* mutants were grown in complete defined medium (SAAC) +/- Arg. Growth curves depict the mean of 3 replicates. Error bars represent 1 standard deviation. *C. sporogenes* is unable to grow in SAAC without arginine (white circles). The growth of wild-type *C. sporogenes* (blue) in SAAC is similar to that of an Δ*adi* mutant (gray), but growth is substantially impaired by deletion of *otc* (green). **(B)** Intracellular ATP levels in wild-type *C. sporogenes* and Δ*adi* and Δ*otc* mutants incubated in PBS containing 2 mM Arg are shown. Wild-type *C. sporogenes* and Δ*adi* generate ATP in SAAC, whereas Δ*otc* does not.

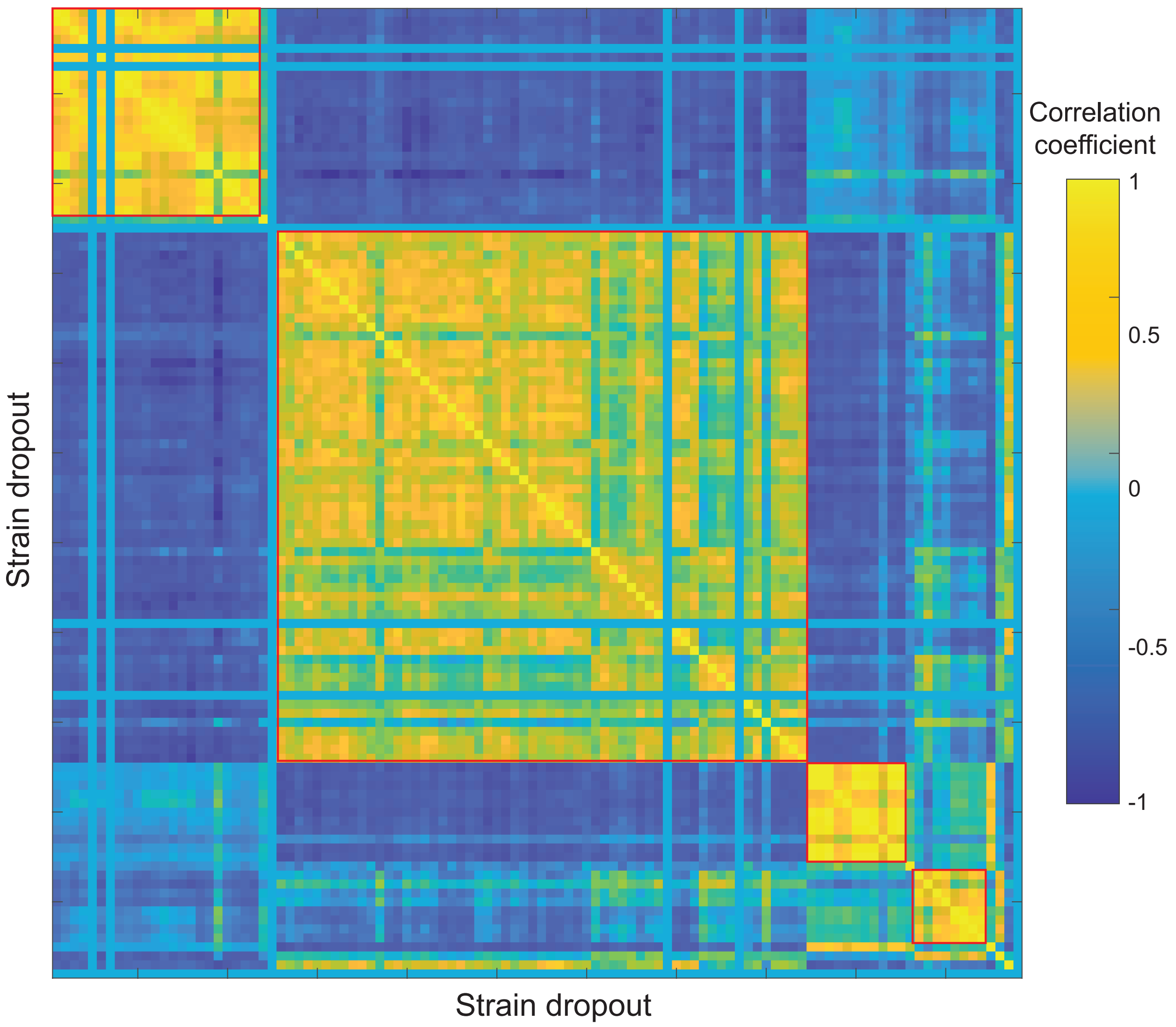


**Figure S6: Relative abundance *z*-score profiles are correlated across strain dropout samples, revealing batch effects resulting from community construction.** Strains are numbered identically on the *x*- and *y*-axes. Four clusters (outlined in red) emerged, all of which correspond to the timing of community assembly or derive from neighboring locations on the plate of strain-dropout communities. As a result, we focused on the two largest clusters and analyzed them separately.

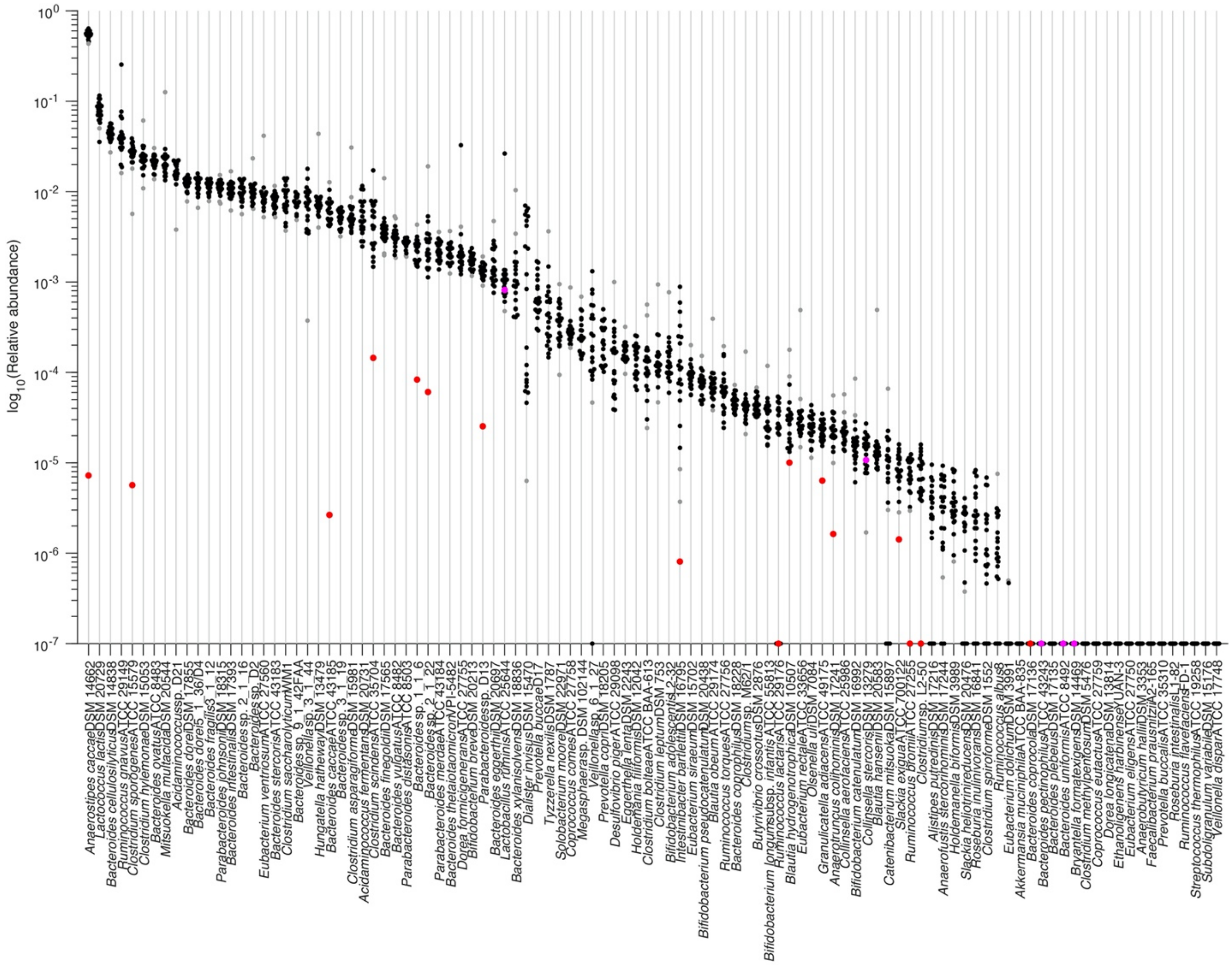


**Figure S7: Strain relative abundances are generally tightly distributed across strain dropouts.** Each column depicts the relative abundance of an individual strain across a set of 24 strain dropouts (black dots), the second largest cluster from **Figure S6**; the relative abundance of each strain in its own dropout is shown as a red dot. Relative abundances in samples with |*z*|>2 are shown as gray dots. Undetected strains were set to $10^{-7}$ for visualization.

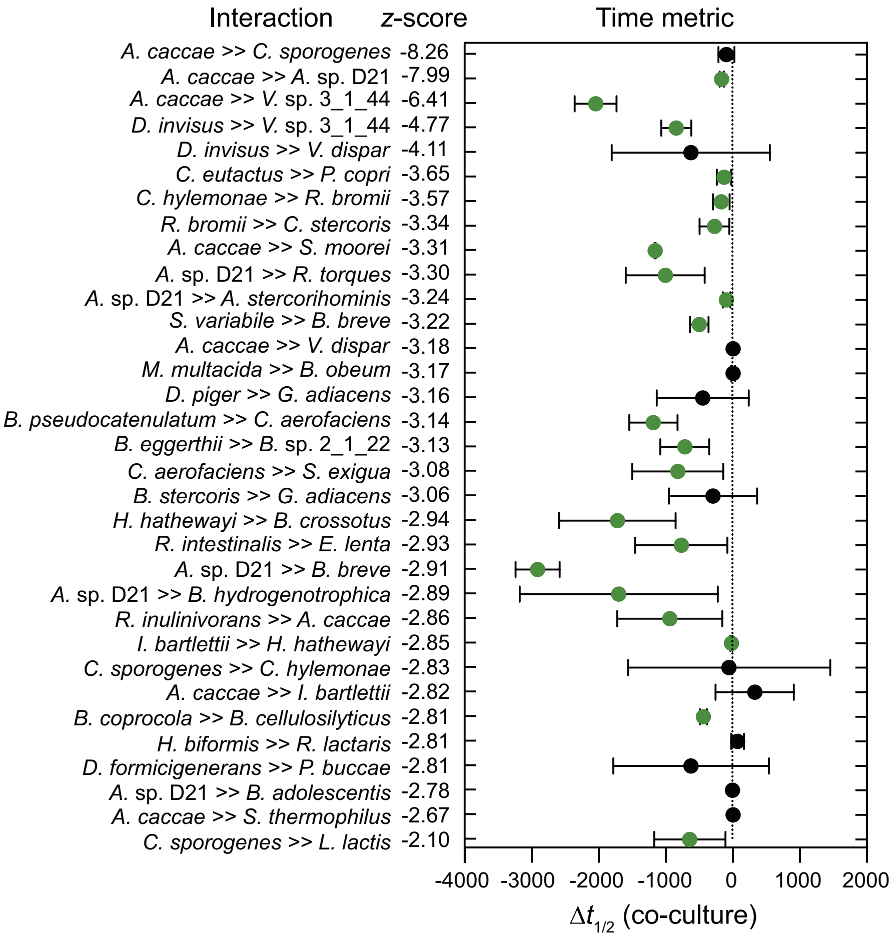


**Figure S8: Growth kinetics of mono-cultures and co-cultures for strains in 32 predicted pairwise interactions with negative *z*-scores**. Removal of the strain on the left was predicted to inhibit the growth of the strain on the right. The *z*-score represents the strength of the interaction. Strains were inoculated individually or as co-cultures in a defined medium (SAAC) and optical density at 600 nm was measured to assess growth. All experiments were performed in technical triplicate with 2-6 biological replicates. For each growth curve, the time to reach half the maximum absorbance ($t_{1/2}$) was computed. The difference between the co-culture $t_{1/2}$ and the minimum of the $t_{1/2}$ values of the individual strains is plotted. Error bars represent 1 standard deviation from the mean. Student's t-test was used to identify significant changes ($p$<0.05). Significant and non-significant changes are shown in green and black, respectively.

### Supplemental Table Legends

**Table S1: Growth medium used for each strain prior to pooling.**

**Table S2: Strain relative abundances in amino acid dropout experiments.**

**Table S3: Strain relative abundances in strain dropout experiments.**

**Table S4: Predicted interspecies interactions based on strain dropout experiments.**

**Table S5: Genome assemblies from strains in the synthetic community.** Each row is a unique genome assembly that was used for all analyses. Columns after the "Source" column indicate the type of assembly and the libraries involved. This procedure resulted in the replacement of eight genomes: two obtained from a PacBio and Illumina hybrid assembly and six from short-read assembly of the respective isolate samples followed by binning.

**Table S6: Modified Standard Amino Acid Complete (SAAC) medium recipe.** The original reagent list for SAAC is based on (Dodd et al., 2017). SAAC complete contains all amino acids at 1 mM concentration, with the exception of cysteine at 4.126 mM.

**Table S7: Strains and primers used in this study.**
